# Histone marks retained during epigenetic reprogramming and their roles essential for fish early development

**DOI:** 10.1101/2022.03.27.486004

**Authors:** Hiroto S. Fukushima, Hiroyuki Takeda, Ryohei Nakamura

**Affiliations:** Department of Biological Sciences, Graduate School of Science, The University of Tokyo, Tokyo, 113-0033, Japan

**Keywords:** Epigenetic reprogramming, histone modification, development, epigenetic memory, fish epigenome, telomere, zygotic genome activation, maternal to zygotic transition

## Abstract

Reprograming of epigenetic modifications after fertilization is required for proper embryonic development and cell differentiation. However, histone modifications that escape reprogramming in non-mammalian vertebrates and their potential functional roles are poorly understood. Here, we quantitatively analyzed histone modification dynamics during reprogramming in Japanese Killifish, medaka (*Oryzias latipes*) embryos, and revealed that H3K27ac, H3K27me3 and H3K9me3 are retained, while H3K4 methylation is completely erased. Furthermore, we experimentally demonstrated the functional roles of such retained modifications at early stages; H3K27ac at promoters is required for proper patterning of H3K4 and H3K27 methylation at zygotic genome activation (ZGA) and specific retention of H3K9me3 at telomeric regions maintains genomic stability during cleavage stage. These results expand the understanding of diversity and conservation of reprogramming in vertebrates and unveil previously uncharacterized functions of histone modifications retained during epigenetic reprogramming.

## Introduction

After fertilization, the epigenetic landscapes of sperm and oocyte chromosomes undergo drastic erasure and re-establishment to ensure totipotency and/or pluripotency in early embryos (Eckersley-Maslin et al., 2018; Xia and Xie, 2020). In mice, DNA methylation and histone modifications transmitted from parental germ cells are globally reset in early embryos. However, some modifications are known to persist and play essential roles in later development (Xia and Xie, 2020). Such examples include DNA methylation at specific gene loci passed on to imprinted genes (Eckersley-Maslin et al., 2018) and the asymmetric pattern of the heterochromatin mark H3K9me3 retained until the blastocyst stage (Wang et al., 2018). Furthermore, a non-canonical pattern of the active mark H3K4me3 established in oocytes is transferred to the zygote and exists until zygotic genome activation (ZGA)(Dahl et al., 2016; Zhang et al., 2016). Additionally, the repressive mark H3K27me3 transmitted from oocytes represses DNA methylation-independent imprinting in embryos (Inoue et al., 2017).

Epigenetic reprogramming of non-mammalian vertebrates is largely distinct from that of mammals. DNA methylation does not undergo genome-wide de-methylation, and is largely maintained (Jiang et al., 2013; Macleod et al., 1999; Potok et al., 2013; Veenstra and Wolffe, 2001). By contrast, histone modifications are more extensively erased in early embryos of non-mammalian vertebrates (Xia and Xie, 2020). Importantly, unlike in mammals, ZGA and subsequent cell differentiation occur around ten cell cycles after fertilization in non-mammalian vertebrates (Jukam et al., 2017), and the reprogramming of histone modifications takes place well before ZGA. For example, in zebrafish and *Xenopus*, H3K4me3, H3K27me3 and H3K9me3 were reported to be almost completely absent before ZGA, but these marks start accumulating at the onset or shortly before the ZGA (Akkers et al., 2009; Hontelez et al., 2015; Laue et al., 2019; Lindeman et al., 2011; Vastenhouw et al., 2010; Zhang et al., 2018a). Thus, parental histone modifications are less likely to be transmitted and maintained through cleavage stages, and to affect later development in non-mammalian vertebrates.

Nevertheless, recent studies suggested that a few types of histone modifications can be inherited or maintained through the cleavage stages to the ZGA in non-mammalian vertebrates. In zebrafish, nucleosomes with H3K4me1, H3K27ac, and H2A.Z (so called “Placeholder nucleosomes”) were suggested to persist after fertilization to pre-mark active and poised promoters (Hickey et al., 2022; Murphy et al., 2018). Additionally, Zhang et al. revealed that H3K27ac accumulates in 4-cell-stage zebrafish embryos and could play a role in priming gene promoters for later ZGA (Zhang et al., 2018a), raising the possibility that histone modifications are maintained before ZGA and required for later development in non-mammalian vertebrates. However, previous studies mostly relied on immunofluorescence staining or conventional chromatin immunoprecipitation-sequencing (ChIP-seq) to demonstrate the presence of histone modifications in early embryos, both of which do not allow quantitative comparison of histone modifications at each genomic locus at different developmental stages (Bonhoure et al., 2014; Li et al., 2014; Orlando et al., 2014). Thus, it remains to be elucidated, exactly to what extent each histone modification is retained during epigenetic reprogramming of non-mammalian vertebrates, and whether the retained modifications affect the phenotype of embryos. Moreover, recent studies revealed that epigenetic patterns are very diverse even in mammals (Lu et al., 2021; Xia et al., 2019), raising the question of what degree the reprogramming process of histone modifications is conserved among vertebrates.

Here, we quantitatively validated the epigenetic reprogramming of histone modifications in Japanese killifish (*Oryzias latipes*, medaka) embryos by conducting quantitative ChIP-seq (Bonhoure et al., 2014; Li et al., 2014; Orlando et al., 2014), together with immunofluorescence staining. We compared the pattern and accumulation levels of histone modifications (H3K27ac, H3K27me3, H3K9me3, H3K4me1, H3K4me2, and H3K4me3) at four developmental stages before and after ZGA. We found that, although all histone modifications were more or less subject to erasure, some histone modifications escaped complete erasure. Importantly, our study suggests that such residual histone modifications are required for maintenance of chromosome stability during cell cleavage and proper establishment of the zygotic epigenetic landscape.

## Results

### Erasure and retention of histone modifications before ZGA in medaka embryos shown by immunofluorescence staining

Like other non-mammalian vertebrates (Jukam et al., 2017), medaka embryos undergo rapid and synchronous cell cycles after fertilization, and cell division gradually becomes longer and asynchronous from the late morula stage (stage 9, 256-512 cells) (Figure 1A and S1). ZGA occurs from the early blastula stage (stage 10, ∼1000 cells)(Nakamura et al., 2021). Blastomeres are pluripotent until the late-blastula stage (stage 11, 2,000-4,000 cells) (Kinoshita et al., 2009), but thereafter cell differentiation begins and three germ layers emerge from the pre-early gastrula stage onwards (stage 12)(Figure 1A).

**Figure 1.**
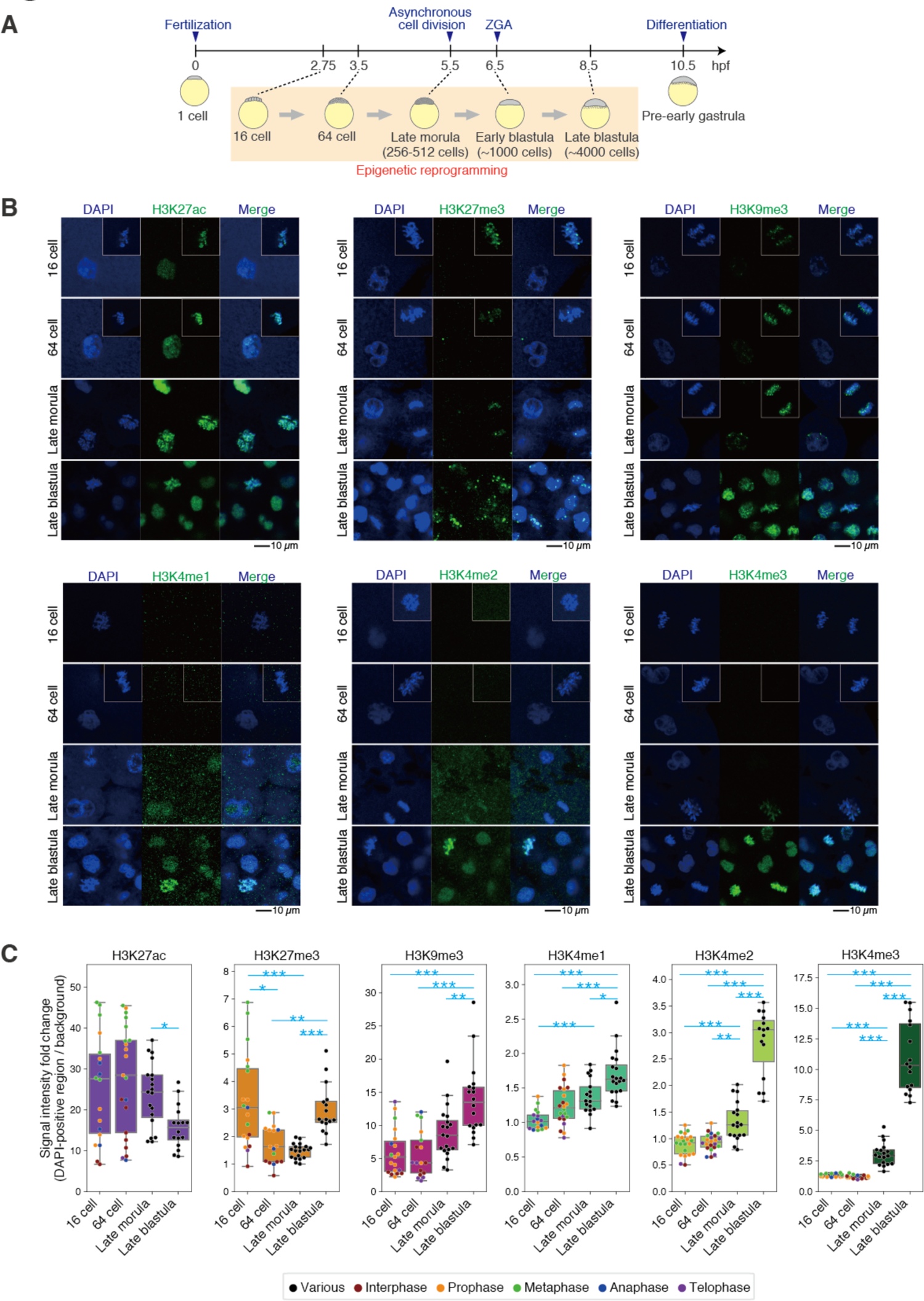
Erasure and retention of histone modifications before ZGA in medaka embryos shown by immunofluorescence staining. (A) A schematic of medaka development. (B) Immunofluorescence staining of medaka embryos at four stages. The nuclei during mitosis from other embryos are inserted in white boxes. The scale bar indicates 10 µm. (C) Boxplots showing the signal intensities of each histone modification in DAPI-positive regions. Each dot indicates the average intensity in a single embryo. The intensity was normalized by background intensity. Cell cycle phases at 16-cell stage and the 64-cell stage embryos are shown as dots with different colors. Phases of cell cycle after the late morula stage are not shown because cells divide asynchronously from the late morula stage. *** p < 0.001, ** p < 0.01, * p < 0.5, respectively.

As a first step toward understanding of when and how histone modifications are reprogrammed in medaka early development, we performed immunofluorescence staining of embryos using antibodies against various histone modifications (H3K27ac, H3K27me3, H3K9me3, H3K4me1, H3K4me2, and H3K4me3) at the 16-cell, 64-cell, late morula and late blastula stages (Figure 1B and 1C). First, we found that among the active marks, H3K27ac was detected at all four stages examined, but other active marks (H3K4me1, H3K4me2 and H3K4me3) were not detected until the late morula stage (Figure 1B and 1C). H3K4 methylation became weakly detectable at the late morula stage and clearly accumulated at the late blastula stage (Figure 1B and 1C). The absence of H3K4 methylation after fertilization is consistent with previous studies in zebrafish (Lindeman et al., 2011; Vastenhouw et al., 2010).

Second, the repressive H3K27me3 and H3K9me3 marks were only detected in mitotic phase chromatin at the 16-cell, 64-cell and late morula stages, but detected in both mitotic phase and interphase chromatin at the late blastula stage (Figure 1B and 1C). This suggests that H3K27me3 and H3K9me3 mostly accumulate during or after ZGA, consistent with previous studies in zebrafish and *Xenopus* (Akkers et al., 2009; Laue et al., 2019; Zhang et al., 2018a), but the two modifications exist at low levels before ZGA, so that they can be detected in compact mitotic chromosome in medaka. We speculated that chromatin compaction during the mitotic phase increased the signal intensity, compared to the dispersed signal observed in interphase nuclei, since the H3K27ac signal in mitotic phases also tended to be higher than that in interphase.

Importantly, the immunofluorescence signals of H3K27ac, H3K27me3 and H3K9me3 were almost completely eliminated by treatment with A485, an inhibitor of H3K27ac histone acetyltransferase CBP/p300 (Hogg et al., 2021; Lasko et al., 2017; Narita et al., 2021; Pelham-Webb et al., 2021) (Figure S16A and S16B), overexpression of a medaka H3K27me3 demethylase Kdm6b (*Oryzias latipes* Kdm6ba, olKdm6ba)(Agger et al., 2007; Inoue et al., 2017; Jullien et al., 2017) (Figure S2A-S2C) and overexpression of a human H3K9me3 demethylase Kdm4d (*Homo sapiens* Kdm4d, hsKdm4d) (Jullien et al., 2017; Matoba et al., 2014; Zhang et al., 2018b) (Figure 4A, 4B and S11A), respectively. These results supported the conclusion that the immunofluorescence staining signals of retained histone modifications were specific. In summary, these data suggests that histone modifications are largely and globally erased before ZGA, but some histone modifications (H3K27ac, H3K27me3 and H3K9me3) might escape from complete erasure.

### Quantitative ChIP-seq reveals both complete erasure and retention of histone modifications during reprogramming in medaka embryos

Immunofluorescence staining is not only non-quantitative, especially in the case of different nucleus size as observed in medaka embryos (e.g. the 16-cell stage versus the late blastula stage) (Figure S1B and 1B), but also does not provide reprogramming dynamics at each genomic region. For this reason, conventional ChIP-seq is frequently used to examine the enrichment of histone modifications at specific genomic loci, but it does not allow comparison of the accumulation levels of modifications between samples due to the lack of normalization of samples (Bonhoure et al., 2014), e.g. among different developmental stages (Li et al., 2014). We thus performed quantitative ChIP-seq or “spike-in” ChIP-seq (Bonhoure et al., 2014; Li et al., 2014; Orlando et al., 2014) to investigate the relative amount of each histone modification along the genome among different developmental stages in medaka. We prepared internal reference (or spike-in) chromatin from zebrafish fibroblast cells (BRF41) and mixed it with experimental chromatin (medaka embryo chromatin) in a same tube, which was then subjected to ChIP-seq (Figure S3). Sequenced reads were aligned to the medaka and zebrafish concatenated genome (Figure S3). Theoretically, the accumulation levels of modifications in the reference chromatin should be the same in all samples, so that the level of histone modifications in medaka chromatin (experimental chromatin) could be normalized using the zebrafish chromatin (reference chromatin) ChIP signal (Figure S3, see also Method). We confirmed that the reference chromatin ChIP-seq pattern was reproducible for all samples (Figure S4), and spike-in normalization improved the reproducibility of experimental chromatin between two biological replicates, as previously reported (Bonhoure et al., 2014) (Figure S5A). We noted that sometimes the correlation between replicates was relatively low (Figure S5A, H3K4 methylations at the 16-cell and 64-cell stages), but that those samples were derived from stages when histone modifications were poorly detected by immunofluorescence staining. In addition, the results of late blastula embryos were very similar to our published data (Nakamura et al., 2014) (Figure S5B). Furthermore, the distribution tendency of spike-in ChIP-seq peaks in the medaka genome is largely consistent with that in other vertebrates (Barski et al., 2007; Zhou et al., 2011) (Figure S5C). For example, three quarters of the H3K4me2 and H3K4me3 peaks are located in promoters, while the heterochromatin mark H3K9me3 tends to be enriched in gene bodies and intergenic regions (Figure S5C). These data indicate that our “spike-in” ChIP-seq signals are reproducible and specific.

We first confirmed that dynamics of global histone modification levels in medaka embryos calculated using spike-in ChIP-seq data (see Method) were similar to those obtained by immunofluorescence staining (Figure S6A). We then asked how the pattern of each modification in the genome is reprogrammed by calculating normalized enrichment levels (termed RPKMspike) (see Method) (Figure 2A and 2B). H3K27ac accumulation was detected and showed similar patterns at all stages examined, but its levels decreased from the 16-cell to 64-cell stages, followed by gradual accumulation onward, indicating that both retention, erasure and addition of these modifications indeed take place (Figure 2A and 2B). H3K27me3 also showed dynamics similar to H3K27ac, but erasure was more intensive (Figure 2A and 2B). On the other hands, H3K4 methylation was almost undetectable before the late morula, suggesting its complete clearance (Figure 2A and 2B). H3K9me3, the heterochromatin mark, also showed global erasure for most regions but was retained at specific genomic regions (described later) (Figure 2A and 2B). We note a slight difference between immunofluorescence staining and spike-in ChIP-seq: H3K4me1 accumulation was low but detected by ChIP-seq at the 16-cell stage (Figure 2A and 2B), while not by immunofluorescence staining (Figure 1A and 1B). This could be due to the technical limitation of immunofluorescence staining against samples with different nuclear size. In summary, spike-in ChIP-seq revealed that, all histone modifications undergo erasure at early stages, albeit to varying degrees, and that H3K27ac, H3K27me3 and H3K9me3 are not completely cleared during epigenetic reprogramming.

**Figure 2.**
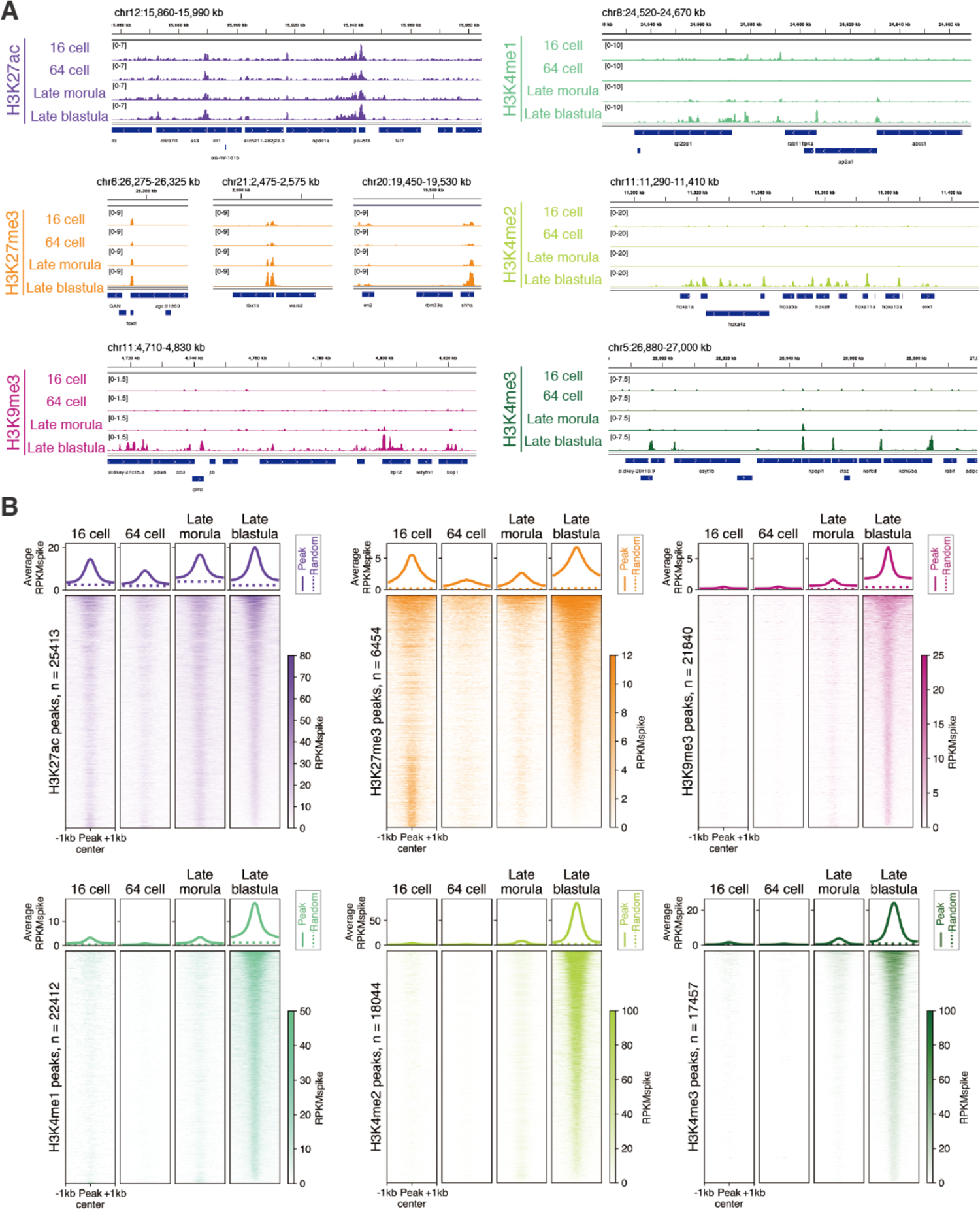
Erasure and retention of histone modifications in early stage medaka embryos quantitatively revealed by spike-in ChIP-seq. (A) Track views showing representative patterns of each modification. Enrichment levels after spike-in normalization are shown. (B) Genome-wide changes in enrichment of each modification. The average enrichment levels after spike-in normalization (RPKMspike) around all peaks and randomized peaks at each stage are shown as solid lines and dashed lines, respectively (top). Heatmaps showing enrichment levels (RPKMspike) around all peaks (± 1 kb from peak center).

It was reported that H3K4me3 primes gene promoters before ZGA in zebrafish (Lindeman et al., 2011). Similarly, our data before spike-in normalization showed the accumulation of H3K4me3 at some promoters in medaka before ZGA (at the late morula stage) (Figure S7A). However, the accumulation levels of those modifications before ZGA were found to be much lower than that after ZGA (at the late blastula stage) by spike-in normalization (Figure S7A and S7B), suggesting that priming of H3K4 methylation before ZGA is rare event in medaka. Intriguingly, only H3K27ac showed comparable levels of accumulation before and after ZGA (Figure 2A and 2B). Therefore, these data indicate the importance of quantitative normalization of ChIP-seq and raise the possibility of a specific role of H3K27ac retention.

### Expression of writers and erasers of histone modifications

The two different dynamics of histone modification reprogramming—complete erasure and residual retention—led us to ask what molecular mechanism underlies this difference. In human embryos, H3K27me3 is globally lost after fertilization, but the same does not occur in mouse embryos. This is thought to correlate with the fact that Polycomb repressive complex 2 (PRC2, a writer and reader complex of H3K27me3) core components, EED and SUZ12, are absent in pre-ZGA human embryos, while present in pre-ZGA mouse embryos (Xia et al., 2019). We thus re-analyzed the expression levels of writers, erasers and related proteins of all histone modifications using our published RNA-seq data (Ichikawa et al., 2017; Nakamura et al., 2021), and confirmed their presence (at least, mRNAs) at all stages examined (Figure S8A-E). Therefore, the presence of mRNAs cannot simply explain the lack or abundance of histone modifications in medaka embryos although we do not know the levels of their translated and activated proteins.

### H3K9me3 at telomeric regions escapes reprogramming

Although the above spike-in ChIP-seq analysis showed extensive erasure of H3K9me3 at most genomic regions (Figure 2A, 2B), immunofluorescence staining detected weak H3K9me3 signals in early-stage embryos (Figure 1B, 1C). We thus investigated in what specific regions of the genome H3K9me3 was accumulating at early stages. Closer scrutiny of ChIP-seq tracks revealed that H3K9me3 was broadly and moderately enriched at chromosome ends at all embryonic stages (Figure 3A, 3B, S9A), although these regions were not called by the conventional peak calling method. This enrichment was observed in most medaka chromosomes (Figure S9B). In vertebrates, telomeres are located at the ends of chromosomes, and characterized by the absence of genes and enrichment of TTAGGG repeats. The regions adjacent of telomeres are called subtelomeres and characterized by low gene density and enrichment of repetitive sequences (Blasco, 2007). Since H3K9me3 is known to accumulate at both telomeres and subtelomeres in somatic cell lines in mouse (García-Cao et al., 2004; Gonzalo et al., 2006), and mainly at subtelomeres in human fibroblasts (Rosenfeld et al., 2009), we hypothesized that H3K9me3 detected at subtelomeres and/or telomeres exceptionally escaped erasure in early-stage medaka embryos.

**Figure 3.**
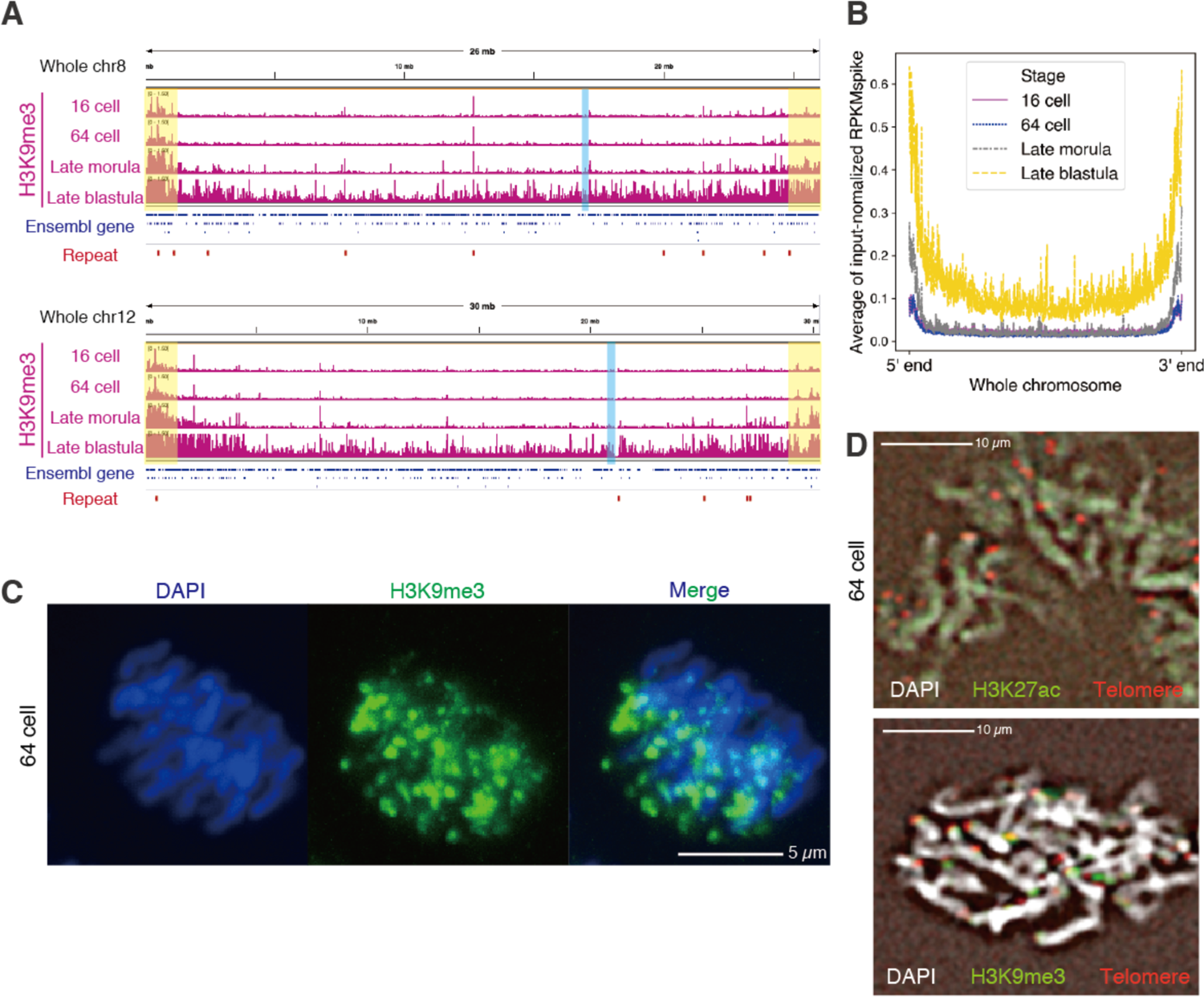
H3K9me3 at telomeric regions escapes reprogramming. (A) Track views showing H3K9me3 accumulation on representative chromosomes. Whole chromosome 8 and chromosome 12 are shown. Ensemble genes are indicated as blue bars, and putative repeat regions (see Method) are indicated as red bars. Chromosome ends are highlighted in yellow. Centromere positions are highlighted in blue. (B) Average H3K9me3 signals on all chromosomes at four stages. To exclude the repeat bias, the signal was subjected to spike-in normalization and further normalized by input signal (see Method). (C) Immunofluorescence staining of H3K9me3 during the mitotic phase (anaphase) at the 64-cell stage. (D) FISH and immunofluorescence staining of telomere repeats and H3K9me3, respectively, using chromosome spreads from embryos at the 64-cell stage.

During the mitotic phase, telomeres are found at the ends of chromosome arms of separated chromatids (Ohta et al., 2011). Thus, we re-analyzed in detail the immunofluorescence data of H3K9me3 staining at the mitotic phase, and found that the H3K9me3 signals tended to locate at chromosome ends during anaphase (Figure 3C and S10A). To further confirm the specific localization of H3K9me3 at telomeres and/or subtelomeres, we examined the colocalization of telomere repeats and H3K9me3 signals. For this, we performed fluorescent *in situ* hybridization (FISH) using probes targeting telomere repeats, followed by immunofluorescence staining of histone modifications, using mitotic chromosomes from 64-cell-stage medaka embryos. As expected, while H3K27ac signal was globally distributed over whole chromosomes, H3K9me3 signal overlapped with telomeres (Figure 3D and S10B). Since only a few chromosomes have telomere repeats in the current medaka genome assembly (Ichikawa et al., 2017), the H3K9me3-enriched regions at chromosome ends detected by ChIP-seq are likely to be subtelomeres. However, due to the limitation of resolution (Figure 3D), we can not further distinguish telomeres and subtelomeres, and hereafter we simply referred them to as “telomeric regions”. Pericentromeres are also known to exhbit H3K9me3 deposition (Dunleavy et al., 2005). However, in medaka, H3K9me3 did not appear to accumulate at pericentromeres at early stages (Figure 3A). Taken together, these observations demonstrate that H3K9me3 escapes reprogramming in early medaka embryos exclusively at telomeric regions.

### Removal of H3K9me3 increases genomic instability before ZGA

Next, we addressed the biological significance of H3K9me3 retention at telomeric regions during reprograming of medaka embryos. We depleted H3K9me3 from early-stage embryos by overexpression of human H3K9me3 demethylase Kdm4d (hsKdm4d) and examined its effect (Figure 4A). We first confirmed that overexpression of hsKdm4d greatly reduced the H3K9me3 signals from the 64-cell stage to the late blastula stage (Figure 4B and S11A). Injected embryos showed impaired gastrulation and severe malformation (Figure S11B and S11C). Strikingly, we found at earlier stages that H3K9me3 depletion increased the proportion of embryos that exhibited abnormal chromosome segregation represented by misaligned chromosomes, lagging chromosomes, DNA-bridge formation, and fusion of nuclei, while control embryos without injection or injected with a catalytically inactive mutant hsKdm4d(H192A) did not show such tendency (Figure 4C, 4D and S11D). Importantly, the increased rate in chromosome-segregation error was observed even at the 64-cell stage, when H3K9me3 accumulates only at telomeric regions in normal embryos (Figure 3A-3D). These data suggest that the residual H3K9me3 at telomeric regions is required for maintenance of chromosome stability during cleavage stages.

**Figure 4.**
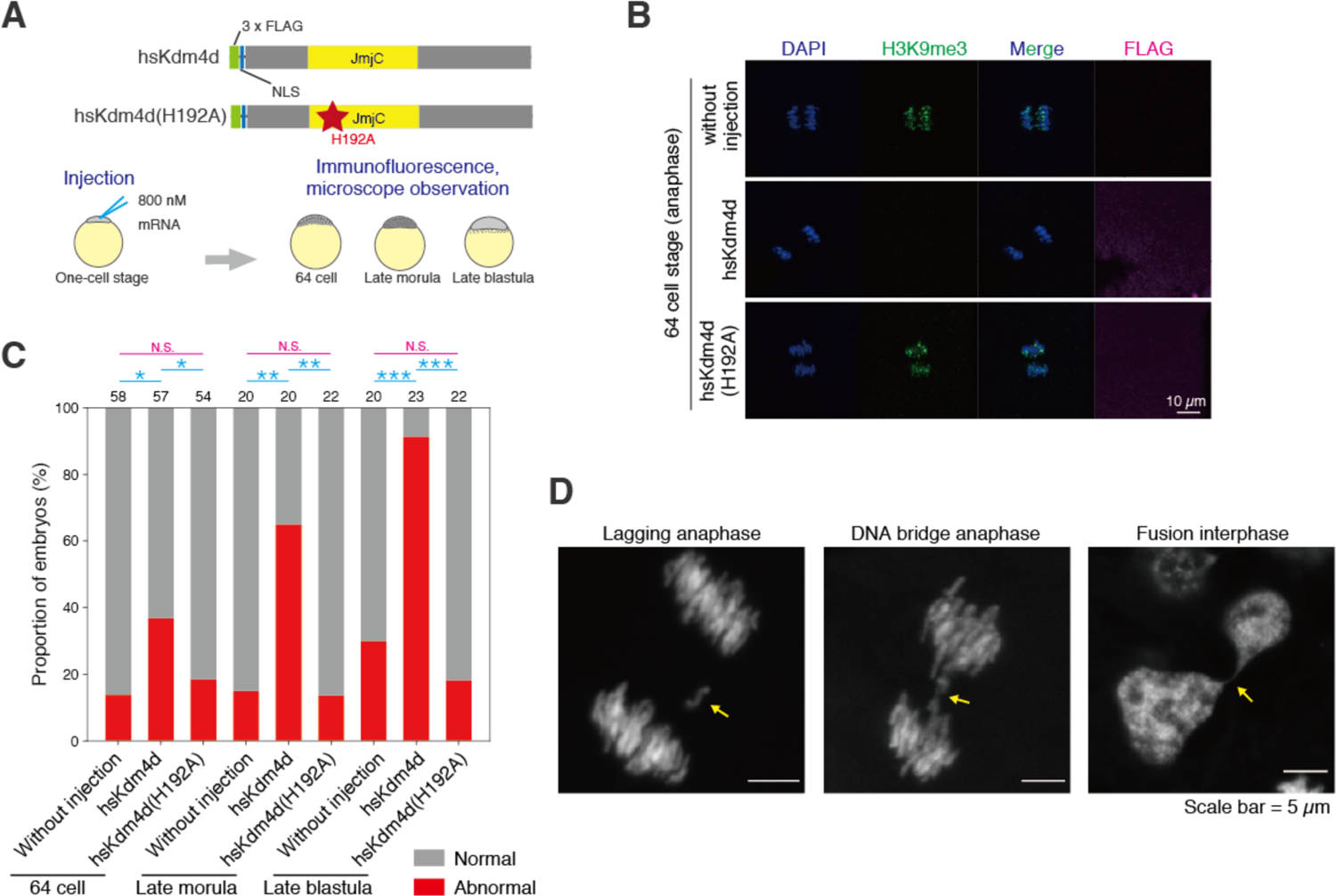
Removal of H3K9me3 increases genomic instability before ZGA. (A) Schematic illustrations of hsKdm4d constructs (wildtype hsKmd4d and its catalytically inactive point mutant hsKdm4d(H192A)) and experimental design. (B) Immunofluorescence staining of H3K9me3 during the mitotic phase (anaphase) at the 64-cell stage after injection of hsKdm4d or hsKdm4d(H192A). (C) Percentage of embryos having abnormal chromosome segregation phenotypes. The number above each bar shows the number of embryos examined in each condition. *** p < 0.001, ** p < 0.01, * p < 0.5, respectively. (D) Representative phenotypes of chromosome segregation errors. Arrows indicates the error.

### H3K27ac pre-marks active and poised promoters

In contrast to the restricted accumulation of H3K9me3 at telomeric regions, H3K27ac marks were globally deposited at promoters and enhancers at all stages examined (Figure 2A and S12A). To investigate the developmental changes in H3K27ac accumulation, we first grouped H3K27ac-marked promoters (n=12,015) into five clusters (Figure 5A-C, S12B-C, see Method). In cluster 1, 2, and 3, H3K27ac was accumulated at the 16-cell stage, then its levels decreased at the 64-cell stage, and increased again from the late morula stage onwards (Figure 5A-C). The dynamics were similar to each other, but the enrichment level was different among these three clusters: cluster 1 - “pre-marked, high”, cluster 2 - “pre-marked, middle” and cluster 3 - “pre-marked, low”, according to their levels at the 16-cell stage. Cluster 4 also exhibits a similar tendency until the late morula stage, but H3K27ac decreased again at the late blastula stage (termed “pre-marked, decreasing”) (Figure 5A-C). Only cluster 5 promoters did not possess H3K27ac at early stages, and H3K27ac started to accumulate from the late morula stage (“Late H3K27 acetylation”) (Figure 5A-C).

**Figure 5.**
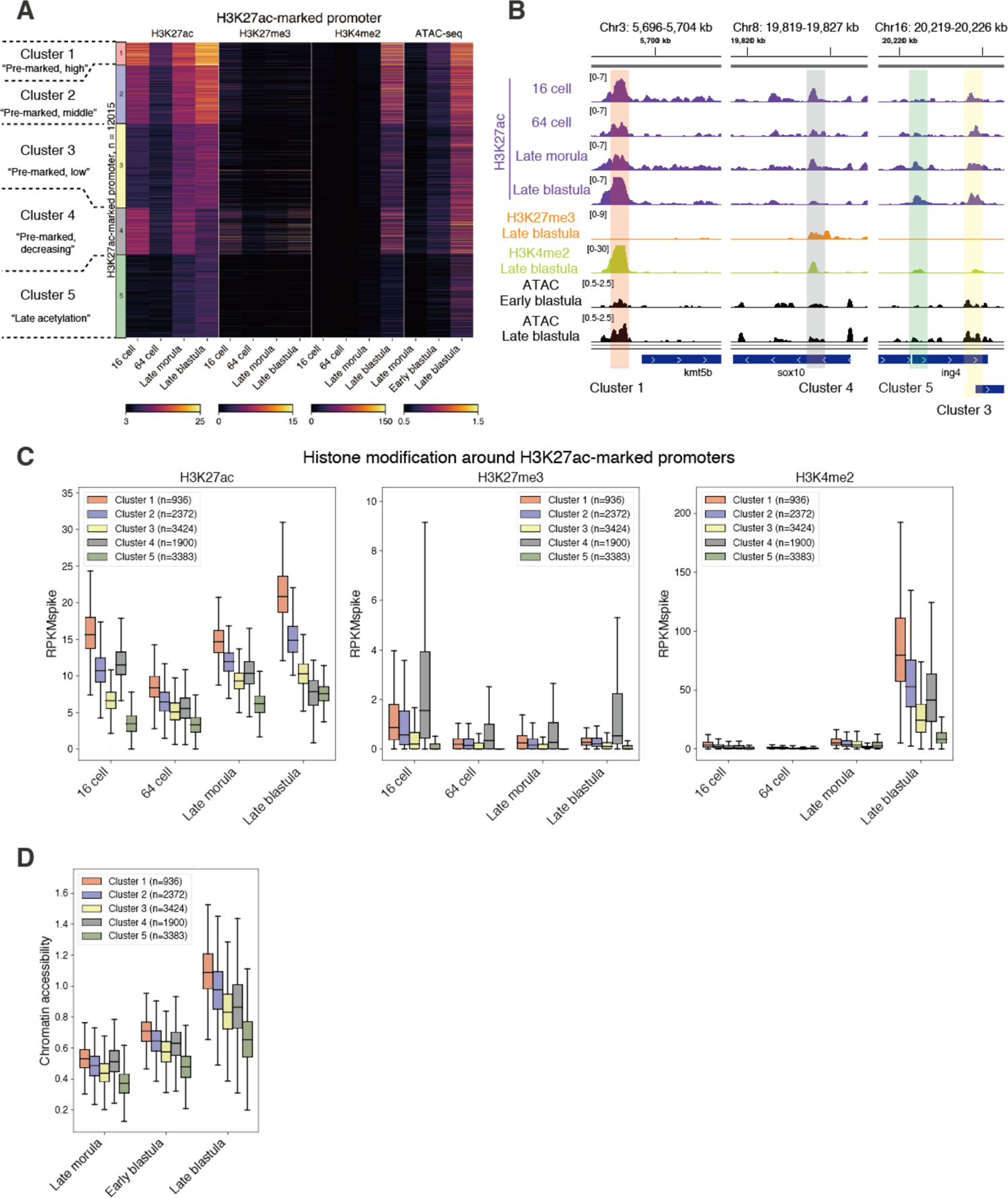
H3K27ac pre-marking is associated with future gain of histone modifications and accessibility at the later stage in medaka. (A) Heatmap showing histone modification enrichments (H3K27ac, H3K27me3 and H3K4me2) and chromatin accessibility (Nakamura et al., 2021) at H3K27ac-marked promoter peaks after clustering. (B) Track views of examples of H3K27ac-marked promoter peaks. The histone modification enrichments after spike-in normalization and chromatin accessibility are shown. Peaks from different clusters are highlighted using different colors. (C) Boxplots showing histone modification levels after spike-in normalization (RPKMspike) in each H3K27ac-marked promoter cluster. The outliers are not shown. (D) Boxplots showing chromatin accessibility around H3K27ac-marked promoters in each cluster.

Cluster 4 was unique in that H3K27me3 accumulation levels increased at the late blastula stage while H3K27ac decreased (Figure 5A-C, S13A). H3K4me2 and H3K4me3 also accumulated at the promoters in cluster 4 from the late blastula stage, thus they are poised promoters (Zhou et al., 2011). Importantly, the reciprocal change in H3K27ac and H3K27me3 was more evident after the onset of differentiation (at the pre-early gastrula stage) in cluster 4 (Figure S13A). Gene ontology (GO) analysis showed that enrichment of terms in cluster 4 is related to developmental genes and transcription factors, which are known targets of PRC2 (Figure S13C). These findings are consistent with the previously described antagonism between H3K27ac and H3K27me3 (Tie et al., 2009) and the switch from H3K27ac to H3K27me3 at developmental gene promoters upon ZGA in zebrafish (Zhang et al., 2018a). Taken together, these data suggest that, in cluster 4, the induction of silencing mediated by H3K27me3 proceeds after ZGA.

### H3K27ac is required for proper accumulation of H3K4 methylation in the late blastula embryos

We then addressed the functional role of H3K27ac pre-marking at promoters in early-stage embryos. Importantly, H3K27ac pre-marking is well correlated with the accumulation of H3K4me2 at the late blastula, i.e. just after ZGA (Figure 5A-C, S13A, and S13B). Moreover, the analysis of our published ATAC-seq data (assay for transposase accessible chromatin followed by high-throughput sequencing) (Nakamura et al., 2021) showed that the promoters pre-marked by H3K27ac (cluster 1-4) become accessible earlier than those without H3K27ac pre-marking (cluster 5); the promoters of cluster 1-4 tended to be open at the early blastula and those of cluster 5 become accessible at the late blastula stage (Figure 5A, 5B, 5D, and S13B). As described above, the switch from H3K27ac to H3K27me3 occurs in cluster 4 promoters (Figure 5A-C, and S13A). Taken together, these observations suggest that H3K27ac pre-marking in promoters at early stages contributes to a later gain of accessibility and subsequent accumulation of H3K4me2 (in cluster 1-4) and H3K27me3 (in cluster 4). As for enhancers, the correlation between H3K27ac pre-marking and rapid gain of H3K4me2 or chromatin accessibility was not so evident (Figure S15A and S15B).

To examine if H3K27ac is required for the later establishment of H3K4me2, H3K27me3 deposition and chromatin accessibility at promoters, we reduced H3K27ac accumulation by treating embryos with the CBP/p300 catalytic inhibitor A485. When embryos were incubated with A485 from the 2-cell stage, the H3K27ac level was globally reduced as shown by immunofluorescence staining (Figure S16A and S16B). This treatment drastically impaired gastrulation (Figure S16C and S16D), consistent with a previous study showing gastrulation arrest after global H3K27ac reduction in zebrafish (Chan et al., 2019).

The effects of H3K27ac reduction were examined by spike-in ChIP-seq using DMSO or A485-treated late blastula embryos. After validating the reproducibility of experiments (Figure S17A), we confirmed that the global H3K27ac level was significantly reduced after A485 treatment (Figure 6A, 6B and S17B). We first compared the H3K4me2 levels at H3K27ac peaks and found that in some H3K27ac peaks, the H3K4me2 level strongly decreased after A485 treatment (“H3K4me2 down”, n=183), but not or weakly affected in other H3K27ac peaks (“H3K4me2 non-sensitive”, n=6903) (Figure 6A and 6B). These “H3K4me2 non-sensitive” peaks had higher levels of H3K27ac than “H3K4me2 down” peaks in A485-treated embryos (Figure 6A-6B) so that residual H3K27 tended to remain even after A485 treatment at “H3K4me2 non-sensitive” peaks. We found that the residual amount of H3K27ac after A485 treatment was correlated with a reduction in H3K4me2 accumulation (Figure S17C). Taken together, the results suggest that H3K27ac is required for proper patterning of H3K4me2 at promoters.

**Figure 6.**
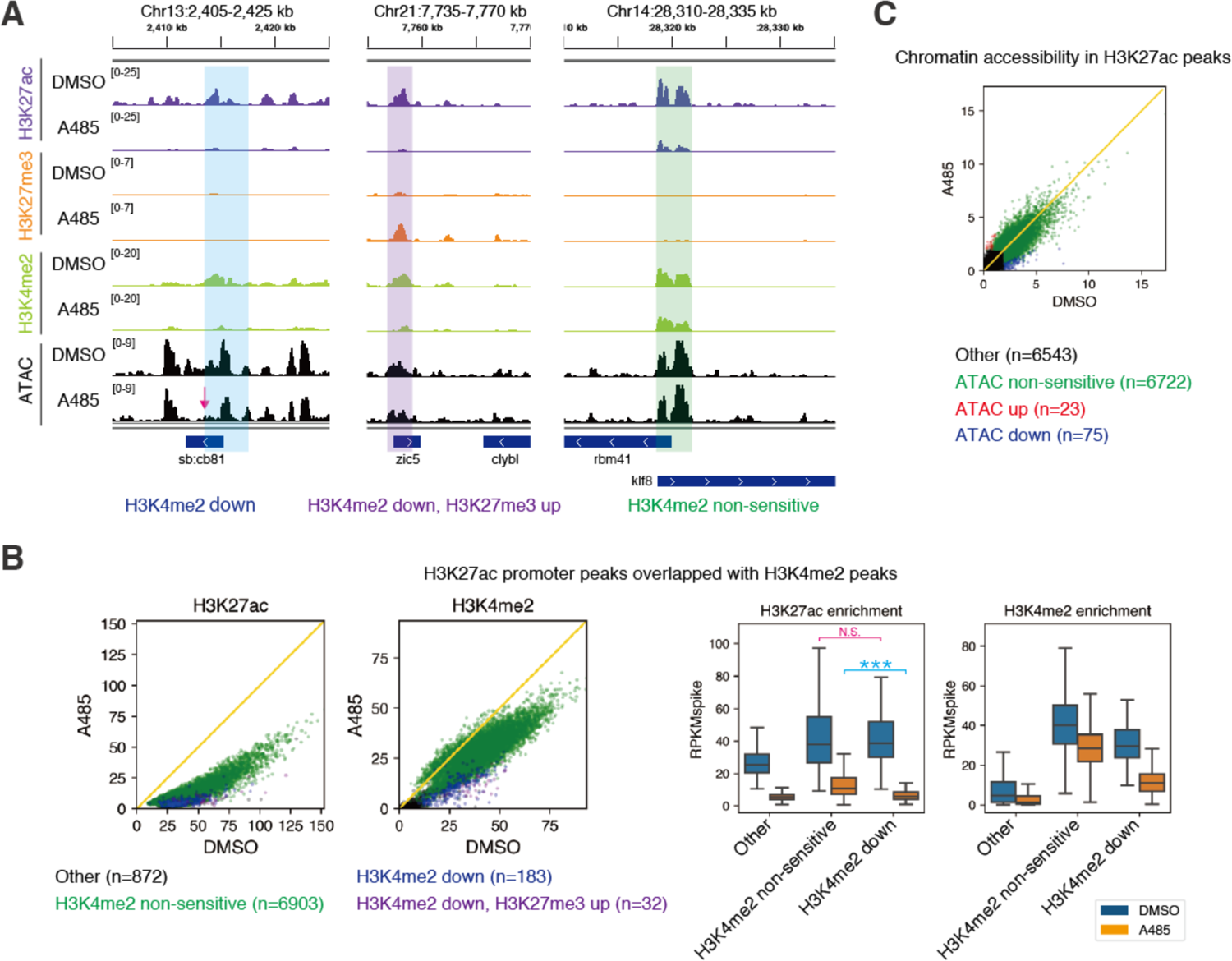
H3K27ac is required for proper histone modification patterning in late blastula embryos. (A) Track views showing the histone modification enrichments after spike-in normalization and chromatin accessibility in DMSO or A485-treated embryos. The magenta arrow indicates where chromatin accessibility was altered after A485 treatment. (B) Comparison of H3K27ac and H3K4me2 enrichment in H3K27ac peaks between DMSO and A485 treatment by scatter plots (left) and boxplots (right). “Other” indicates the H3K27ac peaks without H3K4me2 enrichment. Yellow line in scatter plot indicates y=x. *** p < 0.001. (C) Scatter plot showing chromatin accessibility in H3K27ac peaks in DMSO and A485 treatment. “Other” indicates the H3K27ac peaks with low ATAC-seq signals. Yellow line indicates y=x.

Next, we examined whether H3K27ac affects chromatin accessibility at the late blastula stage by performing ATAC-seq using DMSO or A485-treated late-blastula embryos. After validating the reproducibility and quality of ATAC-seq (Figure S18A and S18B), we found that global reduction of H3K27ac had only minor effects on the ATAC-seq pattern at the late blastula stage (“ATAC non-sensitive” = 6722, “ATAC up” = 23, “ATAC down” = 75) (Figure 6A and 6C). Although some H3K27ac peaks were associated with reduced chromatin accessibility after A485 treatment (Figure 6A and S18C), the number of such changes were very limited (Figure S18D). Consistently, the recent study in zebrafish revealed that H3K27ac is dispensable for chromatin opening during ZGA (Miao et al., 2022). Therefore, we concluded that the gain of chromatin accessibility at the late blastula stage is largely independent of H3K27ac accumulation.

Finally, we examined the effects of H3K27ac on H3K27me3. We found that in most promoters, H3K27me3 accumulation proceeded normally (“Other”: n= 7425, “H3K27me3 non-sensitive”: n=333) in the presence of A485. However, in a small fraction of H3K27ac peaks, reduction of H3K27ac induced abnormal accumulation of H3K27me3 (“H3K27me3 up”, n=232) (Figure 6A and S17D). Thus, in some promoter regions, H3K27ac suppresses the accumulation of H3K27me3 at the late blastula.

For technical reasons, we were unable to deplete H3K27ac specifically and transiently before the late blastula stage, and thus we could not distinguish between the requirement of pre-marking before ZGA or H3K27ac at the late blastula stage. Nevertheless, our data suggest that H3K27ac is required for proper accumulation of H3K4me2 at some promoters, but is dispensable for the gain of chromatin accessibility at the late blastula stage.

### Polycomb-high affinity regions escape complete erasure of H3K27me3 during reprograming

We found weak retention of H3K27me3 at specific sites during reprogramming, although H3K27me3 is extensively erased throughout the genome (Figure 2A and 2B). We classified the H3K27me3 marked regions into three groups (“Erasure”, “Low retention” and “Mild retention”) by the accumulation levels of H3K27me3 at the 64-cell stage, a stage when H3K27me3 enrichment was at the lowest level during reprogramming (Figure S19A). We found that the retained region is associated with the following genetic and epigenetic features of Polycomb high-affinity sequences: (i) High CpG density (Ku et al., 2008; Mendenhall et al., 2010; van Mierlo et al., 2019) and high GCG density (Perino et al., 2018, 2020) (Figure S19B); (ii) DNA hypomethylation (Nakamura et al., 2014) at the blastula stage (Qu et al., 2012) (Figure S19C); (iii) transcription factors and genes related to developmental processes such as Homeobox (Schuettengruber et al., 2017) (Figure S19D-S19F). Interestingly, the analysis of maternally derived transcripts showed that the genes with mild retention are more strongly suppressed in oocytes (Figure S19G), suggesting maternal inheritance of H3K27me3 like in mouse (Inoue et al., 2017), in *Drosophila* (Zenk et al., 2017) and in *Arabidopsis thaliana* (Luo et al., 2020).

## Discussion

Reprograming of epigenetic modification is a critical process during early development. However, the dynamics of histone modification reprogramming and the function of retained modifications had not been fully understood in vertebrates. Here, our quantitative analyses revealed that not all histone modifications are subject to complete erasure in teleost embryos. Among the major histone modifications we examined, H3K4 methylation and H3K9me3 are completely erased during reprogramming but H3K27ac, H3K27me3 and telomeric H3K9me3 are not (Figure 7).

**Figure 7.**
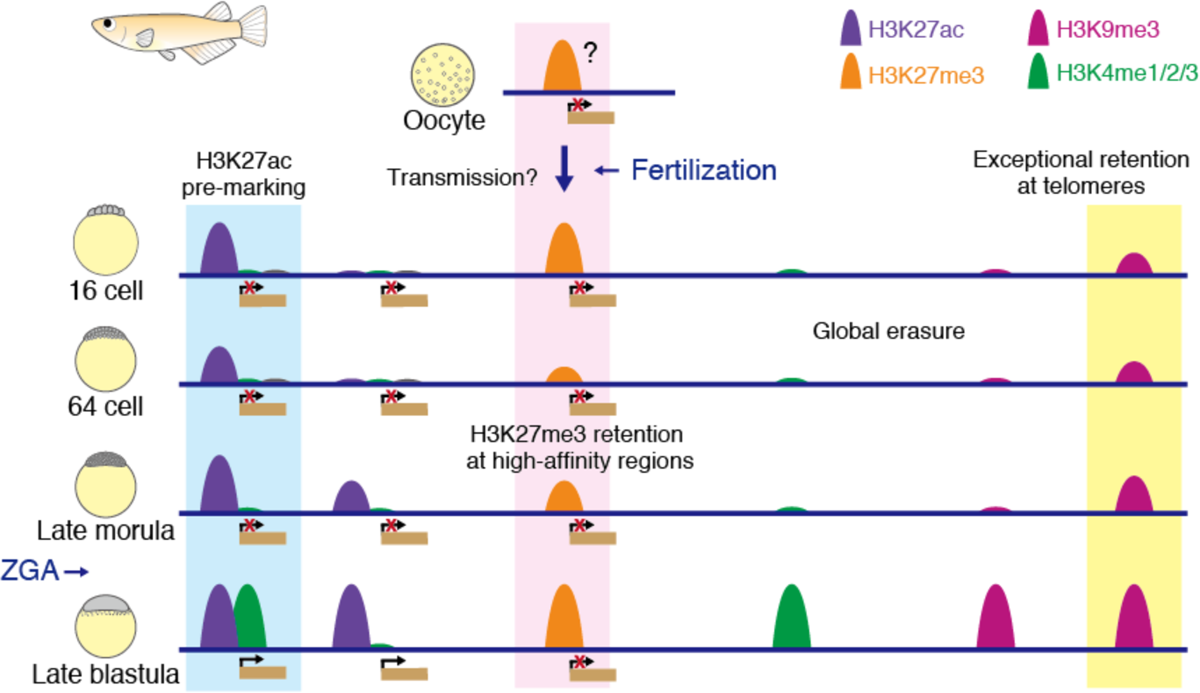
Summary of histone modification reprogramming in medaka. All histone modifications more or less undergo erasure after fertilization, but some modifications escape complete erasure in medaka. H3K27ac pre-marks some promoters and is required for proper patterning of histone modification (e.g. H3K4me2). H3K27me3 is probably transmitted maternally and retained at Polycomb high-affinity regions, although the enrichment was very limited. H3K9me3 at telomeric regions escapes complete erasure and is required for maintenance of chromosome stability during cleavage stage. On the other hand, H3K4 methylations and H3K9me3 except at telomeric regions are extensively erased after fertilization.

### The difference between retained modifications and erased modifications

What mechanism determines whether a modification is retained or erased? At present, these differences cannot simply be explained by the lack or abundance of writers and erasers, as transcripts of all of these are present at all stages (Figure S8).

Because new and free histones are incorporated into chromatin during DNA replication (Stewart-Morgan et al., 2020), the rapid cell cycle in fish early development (Jukam et al., 2017) (Figure 1A) accelerates dilution of all histone modifications. This passive erasure could explain the massive reduction of all histone modifications during reprograming. By contrast, active maintenance should work at least for “retained modifications”. In particular, H3K27ac maintains a comparable level of accumulation during epigenetic reprogramming under rapid cell-cycling (Figure 2A and 2B), and this may require strong *de novo* deposition during cleavage stages. Future investigations are warranted to elucidate molecular mechanisms that enable well-balanced erasure and writing of modifications during early development.

### H3K9me3 protects telomeres

One of the interesting and novel findings of our study is the retention of H3K9me3 at telomeric regions during early development and its role in chromosome stability during cleavage stages. How is abnormal chromosome segregation induced when telomeric H3K9me3 is removed in early embryos? In general, because of the 3’ overhang of telomere repeats, telomeres can be targeted by DNA-damage sensing machinery or DNA repair machinery (O’Sullivan and Karlseder, 2010). To avoid this situation, telomeres are protected by a special protein complex called shelterin, and disruption of shelterin complex results in an increase in genomic instability, which is frequently correlated with cancer development (O’Sullivan and Karlseder, 2010). Notably, the representative phenotypes caused by the failure of telomere protection (telomere dysfunction) are DNA-bridge formation and fusion of two interphase nuclei (Maciejowski et al., 2015; Umbreit et al., 2020), which are similar to those observed in H3K9me3-depleted embryos in this study (Figure 4D and S11D). Thus, our results suggest that the exceptional retention of H3K9me3 at telomeric regions are involved in telomere protection during early development (Figure S20A).

In cultured cells, accumulation of H3K9me3 is broadly observed in heterochromatic regions including telomeres or subtelomeres (García-Cao et al., 2004; Gonzalo et al., 2006; Rosenfeld et al., 2009). A previous study using cultured fibroblast and embryonic stem cells reported that global depletion of H3K9me3, achieved by knocking out H3K9 methyltransferases, resulted in abnormal telomere elongation but did not show telomere dysfunction phenotypes with genome instability observed in our study, such as chromosome fusions (García-Cao et al., 2004). Given that cultured cells are slow-growing, it is possible that cleavage-stage embryos require H3K9me3-dependent telomere protection in addition to shelterin-mediated one to achieve stable chromosomal segregation in rapid cell-cycling.

### The function and molecular mechanisms of H3K27ac pre-marking

Previous studies showed that H3K27ac accumulates from the 4-cell stage in zebrafish embryos (Zhang et al., 2018a), and that inhibition of histone acetyltransferase or acetylation reader disrupted proper ZGA in zebrafish embryos (Chan et al., 2019; Sato et al., 2019; Zhang et al., 2018a). Consistent with these studies, H3K27ac marking is present in medaka embryos before ZGA (Figure 2A and 2B). However, there was no experimental data that explains how H3K27ac contributes to the epigenetic landscape during ZGA. In this study, we experimentally demonstrated that H3K27ac is required for proper accumulation of H3K4me2 at promoters but not for the gain of chromatin accessibility after ZGA (Figure 6A-6C and S17D). Our data is largely consistent with a model previously proposed in zebrafish, based on the correlation of histone modification patterns, that H3K27ac at promoters primes the future accumulation of H3K4me3 (Zhang et al., 2018a).

The molecular mechanism underlying the H3K27ac pre-marking and later gain of H3K4me2 is still unknown. However, it is worth noting that in mammalian cell lines, H3K27ac reader Brd4 was shown to interact with Mediator (Jang et al., 2005), and also that H3K4 methylation writer COMPASS complex is known to interact with Mediator (Quevedo et al., 2019). Furthermore, it was reported that H3K27ac at promoters is recognized by Brd2, another reader of H3K27ac, which in turn mediates H3K4me3 installation and gene activation (Zhao et al., 2021). Thus, induction of H3K4 methylation by H3K27ac in the early medaka embryos may also be mediated by Brd2/4 (Figure S20B).

### Retention of H3K27me3 during reprograming

Residual H3K27me3 was observed in early medaka embryos (Figure S19A), and genes with H3K27me3 retention was strongly repressed in oocytes (Figure S19G). However, the H3K27me3 level at the 64-cell stage (middle of reprogramming) was much lower than that at later gastrulation stage (Figure S6A, S6B). Furthermore, unlike mouse and *Drosophila*, loss of maternally supplied H3K27me3 by maternal Ezh2 knockout did not affect zebrafish development as long as Ezh2 was zygotically expressed (Rougeot et al., 2019). Thus, the biological significance of the retained H3K27me3 for later development in teleosts remains unclear.

In summary, the present study quantitatively determined reprograming dynamics of histone modifications and experimentally demonstrated previously unknown roles of retained histone modifications in early non-mammalian embryos, providing novel insights into epigenetic reprogramming during early embryogenesis.

## Acknowledgments

We acknowledge all the laboratory members for everyday discussion and for their continuous supports. This work was supported by JSPS Grant-in-Aid for JSPS Research Fellow Grant Number JP18J21761 to H.S.F, by Japan Agency for Medical Research and Development (AMED) under Grant Number JP18gm1110007h0001 to H.T., by Japan Society for the Promotion of Science (JSPS) grant number JP21K06013 and Grant-in-Aid for Scientific Research on Innovative Areas grant number JP21H00245 to R.N.

## Author contributions

Conceptualization, Investigation, and Writing – Original Draft, H.S.F.; Writing – Review & Editing, H.S.F, H.T., and R.N.; Supervision, H.T. and R.N.; Funding Acquisition, H.S.F., H.T., and R.N.

## Materials and Methods

### RESOURCE AVAILABILITY

#### Lead Contact

Further information and requests for resources and reagents should be directed to and will be fulfilled by the lead contact, Ryohei Nakamura

### Materials Availability

Materials generated in this study are available from the lead contact upon request.

### Data and Code Availability

Raw reads generated in this study are available at the DDBJ Sequence Read Archive (accession number: PRJDB13204).

### EXPERIMENTAL MODEL AND SUBJECT DETAILS

#### Fish strains and handling

In this study, medaka d-rR strain was used for all experiments. Medaka fish were maintained and raised according to standard protocols (Kinoshita et al., 2009). Fertilized embryos were raised under standard protocols (Kinoshita et al., 2009) at 28°C. Developmental stages were determined based on previously published guidelines (Iwamatsu, 2004). All experimental procedures and animal care were carried out according to the animal ethics committee of the University of Tokyo (Approval No. 20-2).

#### Cell lines

In this study, BRF41 (zebrafish fibroblast cell line) and HEK293 were used. BRF41 was cultured with L-15 medium supplemented with 15% FBS and 1% Penicillin Streptomycin at 33°C. HEK293 was cultured with DMEM.

## METHOD DETAILS

### Immunofluorescence staining and imaging

Fertilized eggs were manually collected and incubated at 28°C. Medaka embryos at appropriate stages were fixed with 4% PFA / PBS for 4 hours at room temperature and overnight at 4°C. After washing and manual dechorionation, embryos were permeabilized with 0.5% Triton-X 100 / 1 x PBS for 30 minutes, washed with PBS for 5 minutes at room temperature 3 times, blocked with blocking buffer (2% BSA, 1% DMSO, 0.2% Triton X-100, 1 x PBS) for 1 hour at room temperature. The primary antibody was added, and samples were incubated at 4°C overnight. Samples were washed with PBSDT (1 x PBS, 1% DMSO, 0.1% Triton X-100) for 15 min 5 times at 4°C, blocked again with blocking buffer at room temperature for 1hr, incubated with blocking buffer including DAPI and secondary antibody for 4 hours at 4°C, washed with PBSDT for 15 min at 4°C 5 times, and finally buffer was replaced by PBS. Blastomeres were isolated from yolk and mounted on slide glasses with 50% glycerol / PBS. Imaging was performed using Zeiss LSM710. All imaging data using the same antibody were taken under same conditions.

### Construction of erasers of histone modifications and their catalytic inactive mutants

Total RNA from two days-post-fertilization medaka embryos and HEK293 were reverse transcribed to cDNA mix using SuperScript III First-Strand Synthesis SuperMix (Invitrogen, 18080400) for cloning of olKdm6ba and hsKdm4d, respectively. olKdm6ba was amplified from the cDNA mix using cloning primers (Forward: GAGAGAAGAAGGACCGCATG, Reverse: TCGGAGAGTCACAGCAGGA). hsKdm4d was amplified from the cDNA mix using cloning primers (Forward: GTACGCTGGTAGATCCTGCT, Reverse: GCATGCCTAGAGTCCTGCAA). PCR products were cloned into the pCRII-TOPO vector (Invitrogen, K465001). The olKdm6ba and hsKdm4d sequences were amplified using primers including 3x FLAG and NLS sequences, and assembled into BamHI and XhoI-linearized pCS2+ vector using NEBuilder HiFi DNA Assembly Master Mix (NEB, E2621). The H377A mutation in olKdm6ba and the H192A mutation in hsKdm4d were induced by PrimeSTAR Mutagenesis Basal Kit (Takara, R046A) using following primer pairs (Forward: GCCGGGTGCCCAGGAGAACAATAATTT, Reverse: TCCTGGGCACCCGGCGTCCGGCTCCC) and (Forward: TGCTTGGGCTACAGAGGACATGGAC, Reverse: TCTGTAGCCCAAGCAAACGTGGTT), respectively. The H377A mutation in olKdm6ba corresponds to a catalytically inactive point mutant of mouse Kdm6b (Kdm6b(H1390A)) in previous studies (Inoue et al., 2017; Xiang et al., 2007). The H192A mutation in hsKdm4d corresponds to the catalytically inactive point mutant of mouse Kdm4d (Kdm4d(H189A)) in a previous study (Matoba et al., 2014).

### *in vitro* transcription

The templates of olKdm6ba, olKdm6ba(H377A), hsKdm4d and hsKdm4d(H192A) mRNA for *in vitro* transcription were generated by PCR from pCS2+ vectors using the following primers (Forward: TGACGTAAATGGGCGGTAGG, Reverse: CAGGAAACAGCTATGAC). mRNA was synthesized using mMESSAGE mMACHINE SP6 Transcription Kit. RNeasy Mini Kit was used to purify RNAs.

### RNA Injection

800 ng / µL (400 nM) of olKdm6ba or olKdm6ba(H377A), or 950 ng / µL (800 nM) of hsKdm4d or hsKdm4d(H192A) mRNAs were injected into one-cell stage (stage 2) embryos.

### Spike-in ChIP-seq library preparation and sequencing

ChIP experiment was performed as previously described (Fukushima et al., 2019) with modifications. Fertilized eggs were collected, dechorionated using hatching enzyme, and incubated at 28°C. Embryos at appropriate stages were transferred into ice-cold PBS in 1.5 mL or 2 mL tube, homogenized by pipetting with P1000 tip, centrifuged for 10 minutes at 4°C and the supernatant was removed for deyolking. For zebrafish fibroblast cell preparation, trypsinized cells were collected and washed with PBS. Subsequently, the deyolked medaka embryonic cell pellet or zebrafish fibroblast cell pellet was resuspended with ice-cold PNPP (PBS containing 20 mM sodium butyrate, 1 mM PMSF, and 1 x protease inhibitor), and cells were cross-linked by adding formaldehyde (1% volume per volume final) for 8 minutes at room temperature then quenched by adding glycine (125 mM final). After washing with ice-cold PNPP, cross-linked cells were stored in −80℃ as a dry pellet. The cross-linked cell pellet was lysed with Lysis-ChIP buffer (50 mM Tris-HCl pH 8.0, 10 mM EDTA, 1% SDS, 20 mM sodium butyrate, 1 mM PMSF, 1 x protease inhibitor) and sonicated in a microTUBE AFA Fiber Snap-Cap 6×16mm using Covaris S220 with optimized parameters (Peak Power: 105, Duty Factor: 4.0, cycles per burst: 200, duration: 750 seconds).

To measure the concentration of chromatin and to confirm successful fragmentation, 10 µL of sonicated chromatin solution was treated with RNase A for 1 hour and proteinase K for 2 hours, and DNA was purified by phenol: chloroform: isoamyl alcohol method and ethanol purification. The DNA concentration was measured by Qubit dsDNA HS Assay Kit, and the average of two tube concentrations was used for subsequent spike-in ChIP. Size distribution of fragmented DNA was examined using Agilent 2100 BioAnalyzer and DNA High Sensitivity Kit. 30 µL of Dynabead Protein A and 3 µg antibody were incubated with RIPA buffer (10 mM Tris-HCl pH 8.0, 140 mM NaCl, 1 mM EDTA, 0.5mM EGTA, 1 % Triton X-100, 0.1 % SDS, 0.1 % sodium deoxycholate) overnight at 4°C. 30 ng of experimental chromatin (from medaka embryos) and 15 ng or 30 ng of reference chromatin (from zebrafish fibroblasts) were used for H3K27ac / H3K27me3 / H3K9me3 ChIP or H3K4 methylation ChIP, respectively. Chromatin solution was incubated overnight at 4°C with RIPA-ChIP buffer (RIPA buffer supplemented with 20 mM sodium butyrate, 1 mM PMSF, 1 x protease inhibitor). Samples were washed 3 times with ice-cold RIPA buffer and once with ice-cold ChIP-TE buffer (10 mM Tris-HCl pH 8.0, 10 mM EDTA), and reverse-crosslinked by incubating in a thermomixer (65°C, 800 rpm) overnight with Lysis / NaCl buffer (50 mM Tris-HCl pH 8.0, 10 mM EDTA, 1% SDS, 300 mM NaCl). Samples were placed on a magnetic stand, and the supernatant was transferred into new tubes, treated with RNase A at 37°C for 2 hours, Proteinase K at 55°C for 2 hours, and DNA was purified by phenol: chloroform: isoamyl alcohol method and ethanol purification. The DNA concentration was measured by Qubit dsDNA HS Assay Kit.

ChIP-seq libraries were prepared using KAPA Hyper Prep Kit from 1 ng ChIPed DNA. When the amount of ChIPed DNA was less than 1 ng, ChIP was repeated once more and the two tubes were combined and used for library preparation. All ChIP-seq libraries were sequenced using the Illumina HiSeq1500 system.

### A485 treatment and ATAC-seq

Fertilized eggs were dechorionated and incubated with 20 µM A485 or DMSO from the 2-cell stage to the late blastula stage. These embryos were used for subsequent immunofluorescence staining, or spike-in ChIP-seq as described above. ATAC-seq using A485-treated or DMSO-treated embryos were conducted as previously described (Nakamura et al., 2021) with modification. Four dechorionated embryos at the late-blastula stage were homogenized in ice-cold PBS, centrifuged at 500 g, 4°C for 5 minutes, and the supernatant was removed. After washing with ice-cold PBS, cells resuspended in 50 µL of ice-cold lysis buffer (10 mM Tris-HCl, pH 7.4, 10 mM NaCl, 3 mM MgCl2, and 0.1% Igepal CA-630), centrifuged for 10 min at 500g, and the supernatant was removed. 50 µL of transposition mix (including 25 µL of TD and 2.5 µL of TDE1 from Nextera Sample Preparation kit) was added and the tube was incubated at 37°C for 30 minutes, and the tagmented DNA was purified using a MinElute PCR purification kit. Subsequently, small DNA fragments were size-selected and amplified by two sequential rounds of PCR. First, nine-cycle PCR was performed using indexed primers from a Nextera Index kit and KAPA HiFi HotStart ReadyMix, and the amplified DNA was size-selected to a size of <500 bp using AMPure XP beads. Then, a second five-cycle PCR was performed using the same primer as the first PCR and purified by AMPure XP beads. ATAC-seq libraries were sequenced using the Illumina NovaSeq 6000 system.

### FISH and immunofluorescence of chromosome spreads

Dechorionated embryos were homogenized at the 64-cell stage by pipetting up and down in 1x PBS. After centrifugation, the pellet was dissolved with fixative (1% PFA with 0.2% Triton-X-100, pH 9.2), cells were spread on glass slides, and incubated overnight at 4°C in a humid chamber. After washing, hybridization solution (50% formamide, 5x SSC, 50 µg/mL heparin, 500 µg/mL tRNA, 0.1% Tween 20, 1.35 mg/mL BSA, 0.2 µM TelC-Cy3, pH ∼6) was placed on a slide. After denaturing at 80°C for 10 minutes, slides were placed in a humid chamber and incubated at 37°C overnight. The slides were washed, blocked with blocking buffer (same as for immunofluorescence staining), and incubated with primary antibody at 4°C overnight. After washing, slides were incubated with secondary antibody at 37°C for an hour. Finally, slides were washed, VECTASHIELD Mounting Medium with DAPI was added to slides, and coverslips were placed on slides. Imaging was performed using Keyence microscope BZ-9000.

## QUANTIFICATION AND STATISTICAL ANALYSIS

### Biological replicates

Immunofluorescence experiments and phenotype analyses were repeated at least twice. Two biological replicates were generated for spike-in ChIP-seq and ATAC-seq.

### Statistical analysis

All box and whisker plots in this study indicate the median (center line), the first and third quartiles (bottom and top edge of the boxes, respectively), and 1.5 times the interquartile range (whiskers). Outliers are not plotted except for images of immunofluorescence staining.

For statistical testing of multiple samples, normality and equal variances were tested first by the Shapiro-Wilk test and the Bartlett test, respectively, with p-value = 0.05 using Python library SciPy (Virtanen et al., 2020). In all tests in this study, the null hypothesis of either normality and/or equal variances was rejected. Therefore, we performed Dwass, Steel, Critchlow and Fligner all-pairs comparison test as non-parametric test for statistical significance validation using the Python package scikit_posthocs (Terpilowski, 2019) (*** p < 0.001, ** p < 0.01, * p < 0.5, respectively). We performed a two-sided Fisher’s exact test only for the multiple comparison of proportion of embryos in Figure 4C, and p-values were normalized by the Holm method using a Python library statsmodels (statsmodels.stats.multitest.multipletests, method=”holm”) (*** p < 0.001, ** p < 0.01, * p < 0.5, respectively).

For statistical testing of two samples, we initially performed a Shapiro-Wilk test as described above and the null hypothesis was not rejected in Figure S16B and rejected in Figure 6B. Next, we tested equal variances by F-test (p-value = 0.05) and the null hypothesis was rejected in Figure S16B. Therefore, we performed Welch’s t-test in Figure S16B and Wilcoxon’s rank-sum test in Figure 6B using SciPy (*** p < 0.001, ** p < 0.01, * p < 0.5, respectively).

### Quantification of immunofluorescence staining results

Fiji (Schindelin et al., 2012) was used for immunofluorescence staining quantification. For measurements of chromatin signal intensity, DAPI-positive regions were chosen and the average of intensities inside DAPI-positive regions in a single embryo was measured. For measurements of background signal intensity, a region outside the nuclei was manually chosen, and signal intensity in that region was measured. Finally, the signal intensity fold change (DAPI-positive region / background) was calculated and used for subsequent statistical tests.

### Spike-in ChIP-seq and ATAC-seq data processing

Low quality reads and adapter-derived sequences were trimmed by Trimmomatic (Bolger et al., 2014). Trimmed-reads were aligned to the medaka (HdrR) and zebrafish (Zv9) concatenated genome and medaka HdrR genome using BWA (Li and Durbin, 2010) for ChIP-seq and ATAC-seq, respectively. Alignments with mapping quality smaller than 20 and PCR duplicates were removed by SAMtools (Li et al., 2009). Spike-in ChIP-seq reads aligned to medaka or zebrafish genome were separated for subsequent analysis. For ATAC-seq data, MACS2 (Zhang et al., 2008) was used to generate signals per million reads tracks with following options: --nomodel --extsize 200 --shift −100-g 600000000 -q 0.01 -B -SPMR, and this value in each peak was used for analysis of chromatin accessibility.

### Previous ChIP-seq data processing

Previous ChIP-seq data (Nakamura et al., 2014) were aligned to medaka HdrR genome, and other processing was performed as described for spike-in ChIP-seq data processing.

### Previous ATAC-seq data processing

The previous ATAC-seq data (Nakamura et al., 2021) were downloaded and processed as described above.

### RPKM calculation

We calculated conventional RPKM (RPKMconv) using the following formula:

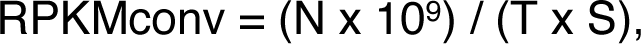

where N is the number of reads mapped in each region, T is the total read number and S is the size (bp) of each region. Also, based on a previous study (Li et al., 2014), we calculated spike-in normalized RPKM (RPKMspike) using the following formula:

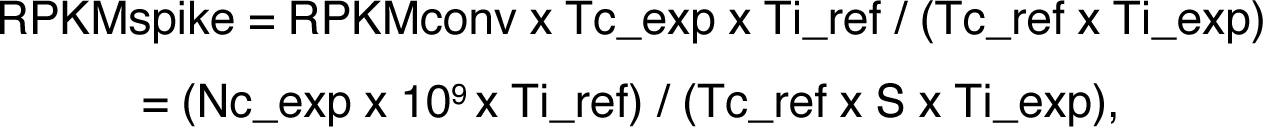

where Nc_exp is number of ChIP reads of experimental chromatin mapped in each region, Tc_exp is number of total ChIP reads of experimental chromatin, Tc_ref is number of total ChIP reads of reference chromatin, Ti_exp is number of total input reads of experimental chromatin, and Ti_ref is number of total input reads of reference chromatin.

### ChIP-seq correlation

To compare spike-in ChIP-seq with conventional ChIP-seq, or between two replicates, the RPKMconv or RPKMspike of each 5 kb window was calculated for the whole genome. Pearson’s correlation between two replicates suggested high reproducibility, so we pooled two replicates for subsequent analysis.

### ATAC-seq correlation

To compare ATAC-seq data with previous data or between two replicates, the RPKMconv of each 5 kb window was calculated for the whole genome. Pearson’s correlation between two replicates suggested high reproducibility.

### Putative repeats calling

To remove regions containing repeat sequences in medaka genome, putative repeat regions were identified as follows: MACS2 (Zhang et al., 2008) peak calling of input data was performed using following parameters: --broad --nomodel --extsize 250 -g 600000000 -q 0.001 -B --SPMR --keep-dup all, called peaks within 500 bp distance were concatenated using bedtools merge (Quinlan and Hall, 2010), and peaks called by more than half of input samples were defined as putative repeat regions.

### ChIP-seq peak calling and generating tracks

MACS2 (Zhang et al., 2008) was used to call peaks and to generate signals per million reads tracks using following parameters: -g 600000000 -q 0.01 -B --SPMR --keep-dup all and -g 1300000000 -q 0.01 -B --SPMR --keep-dup all for experimental chromatin and reference chromatin, respectively. The peaks from the four stages (16 cell, 64 cell, late morula and late blastula) were concatenated if they are within 150 bp using bedtools merge (Quinlan and Hall, 2010), and peaks that overlapped with putative repeat regions defined above were removed. The peaks in unanchored contigs were also removed, and the remaining peaks were used for subsequent analyses. To measure the background enrichment level, random peaks were generated by bedtools shuffle (Quinlan and Hall, 2010) from the merged and survived peaks. For spike-in normalization of track data, the values of normalized pileup track data generated by MACS2 were further normalized as follows:

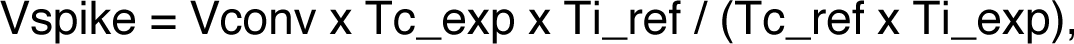

where Vspike is track data value after spike-in normalization, Vconv is original track data value generated by MACS2 --SPMR, and other variables are explained above in the RPKM calculation section. Integrative Genome Viewer (Robinson et al., 2011) was used for visualization of track data before and after spike-in normalization.

### Global histone modification level calculation

Based on a previous paper (Li et al., 2014) with slight modification, the global histone modification level of experimental chromatin (G) was calculated using spike-in ChIP-seq data as follows:

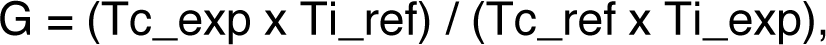

### Peak annotation

All merged peaks that survived after putative repeat removal were associated to nearby genes using Homer annotatePeaks.pl (Heinz et al., 2010). Specifically, peaks within 2.5 kb upstream to 1.5 kb downstream of the transcription start site (TSS) of nearby genes were regarded as peaks in the promoter, and default output of Homer annotatePeaks.pl was used for other annotations (“Exon”, “Intron”, “TTS (transcription termination site)”, “Intergenic”).

### Clustering analysis of H3K27ac peak

We chose peaks within 2.5 kb upstream to 1.5 kb downstream of TSS of nearby genes and peaks in distal regions (at least 2 kb away from promoters) as promoter peaks and enhancer peaks, respectively. If more than two peaks were included in the same promoter, only the representative peak whose RPKMspike was highest was used for promoter peak analysis. RPKMspike value within each peak was used for subsequent clustering. To determine the optimal number of clusters for k-means clustering, we applied the elbow method and the Silhouette method, and we also checked the heatmaps from various numbers of clusters (see also Figure S12 and S14). Finally, we concluded that 5 and 6 is the best number of clusters for promoter and enhancer peak clustering, respectively.

### Classification of H3K27me3 peaks

H3K27me3 peaks were divided into three groups using RPKMspike values of peaks in 64 cell stage (“Erasure”: RPKMspike = 0, “Low retention”: 0 < RPKMspike ≤ 1, “Mild retention”: 1 < RPKMspike). H3K27me3 peaks within 2.5 kb upstream to 1.5 kb downstream of TSS of nearby genes were considered as promoter peaks. If more than two peaks were included in a same promoter, a representative peak whose RPKMspike is highest was used.

### Gene ontology analysis

To find gene ontology terms enriched in promoter peaks, the genes downstream of H3K27ac or H3K27me3 promoter peaks were listed. Gene ontology enrichment analysis was performed by PANTHER (Mi et al., 2019). In Figure S13C, gene ontology terms (biological process) which have FDR less than 10^-5^ and whose enrichment was more than 2.1 fold were listed. In Figure S19F, gene ontology terms (biological process) which have FDR less than 10^-5^ and whose enrichment was more than 3.3 fold were listed. Because many terms were found enriched in the “Mild retention” cluster, the top 20 terms sorted by FDRs were shown in Figure S19F.

### RNA-seq data processing

Previous RNA-seq data (Ichikawa et al., 2017; Nakamura et al., 2021) were downloaded and processed as previously described (Nakamura et al., 2021). Low quality reads and adapter-derived sequences were trimmed by Trimmomatic (Bolger et al., 2014). The reads were aligned to medaka HdrR genome using STAR (Dobin et al., 2013), and alignments with mapping quality smaller than 20 were removed using SAMtools (Li et al., 2009). FPKM or RPKM table was generated by R library DESeq2 (Love et al., 2014).

### Analysis of maternal expression levels

RNAs in 2-cell-stage embryos are maternally derived. Therefore, we used FPKM at the 2-cell stage for analysis of maternal expression levels.

### Whole genome bisulfite-seq data processing

The whole genome bisulfite-seq data from a previous study (Qu et al., 2012) was downloaded from accession number SRA026693. Low quality reads and adapter-derived sequences were trimmed by Trimmomatic (Bolger et al., 2014). Trimmed reads alignment to medaka HdrR genome, deduplication and CpG methylation level calculation were done using bismark (Krueger and Andrews, 2011). Only those CpG dinucleotides with coverage of ≥ 4 were regarded as valid calls and used for subsequent analyses.

### Analysis of DNA methylation level around peaks

DNA methylation level averages in 100 bp window-divided genomic regions around peak center (within peak center ± 2 kb) were calculated and used for heatmap and line plot. If any valid CpG dinucleotides were not included in the 100 bp window-divided genomic region, the average methylation level of that region was accounted to be 100%.

### Analysis of epigenetic changes induced by A485 treatment

Initially, we pooled two biological replicates of ChIP-seq and performed macs2 peak calling as described above. Here, we only chose peaks whose fold change of H3K27ac, H3K27me3 and H3K4me2 enrichment in macs2 peak calling output was higher than 5.5, 7.0 and 10.0, respectively. Those peaks from DMSO- or A485-treatment were next merged as described above. We chose the merged promoter H3K27ac peaks whose fold change (log2(A485/DMSO)) was higher than 0.8 or lower than −0.8 in both replicates. To further discuss the changes in the level of enrichment of other histone modifications, H3K27ac peaks that overlapped with H3K27me3 peaks or H3K4me2 peaks were chosen (the peak center is within peaks of another modification), and reproducible changes of H3K27me3 or H3K4me2 in such peaks were tested as we did for H3K27ac level. Specifically in Figure S17C, the significantly altered H3K27ac peaks that overlapped with H3K4me2 peaks were further divided into “mild down” (y > 0.18 * x) and “severe down” (y ≤ 0.18 * x).

To compare the chromatin accessibility in H3K27ac peaks, ATAC-seq data was processed as described above. The peaks whose accessibility (output value of ATAC-seq MACS2) was higher than 2 at least in one sample and whose fold change of accessibility (log2(A485/DMSO)) was higher than 1.2 or lower than −1.2 were classified as significant changes.

### KEY RESOURCES TABLE

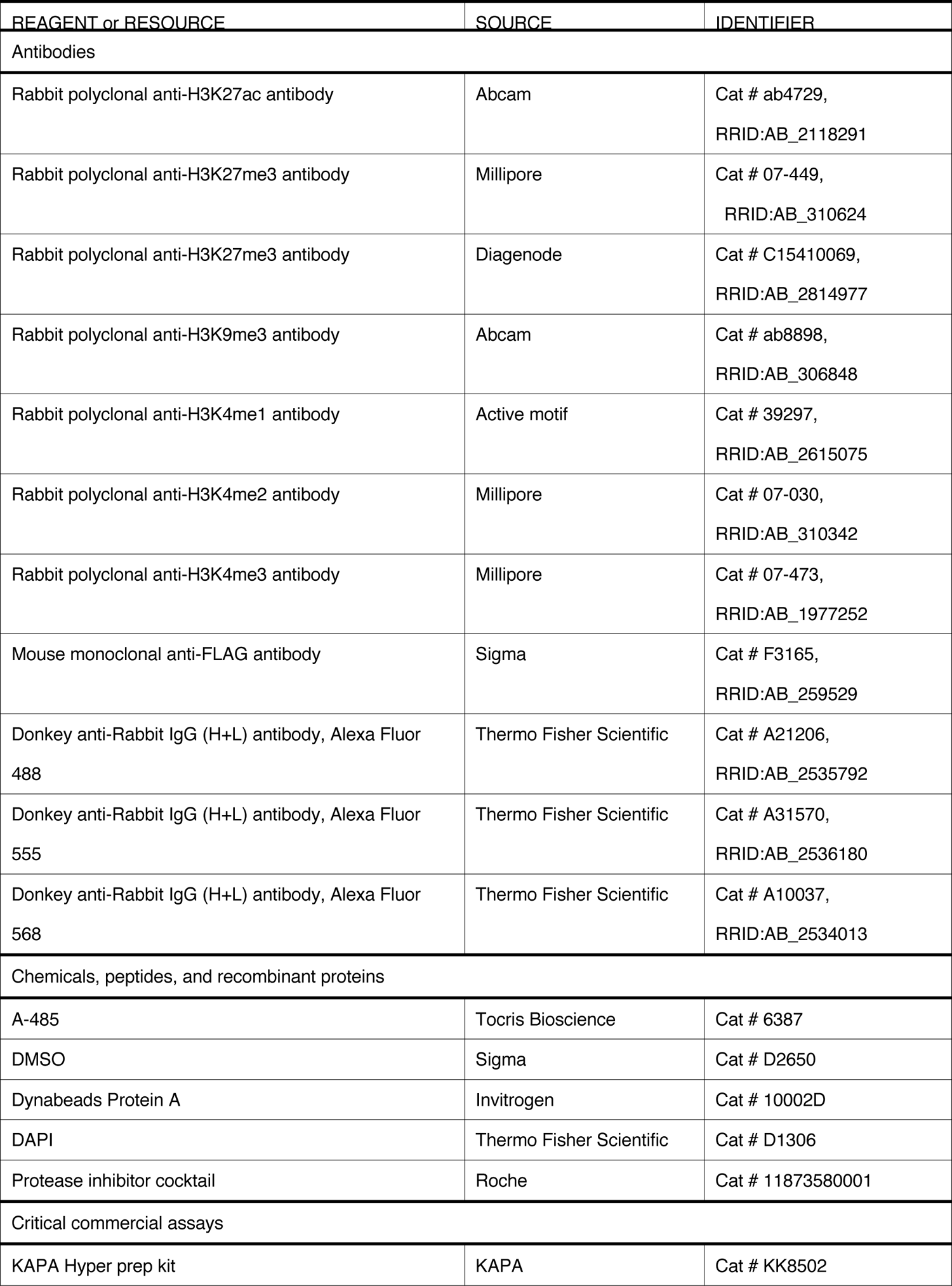
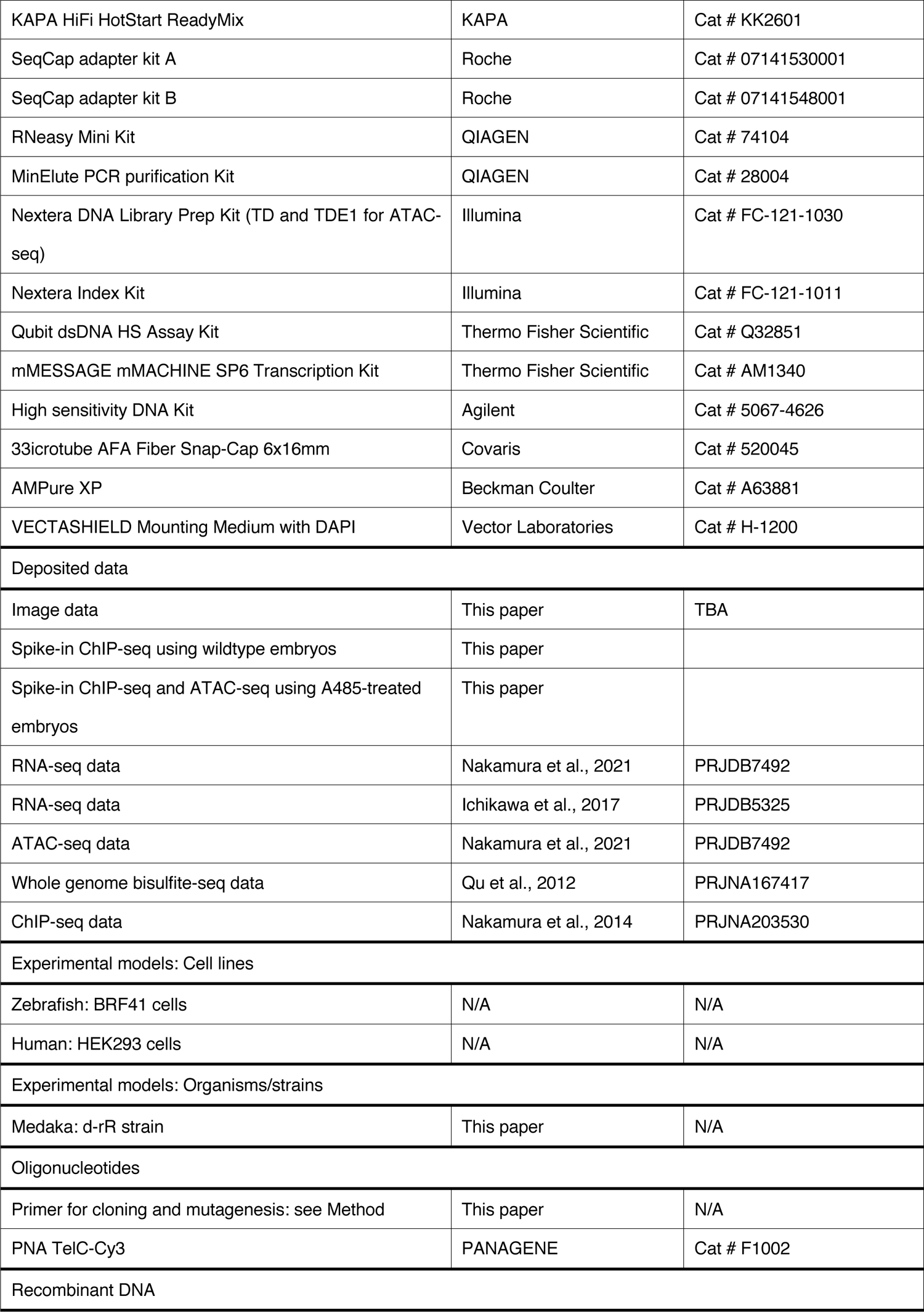
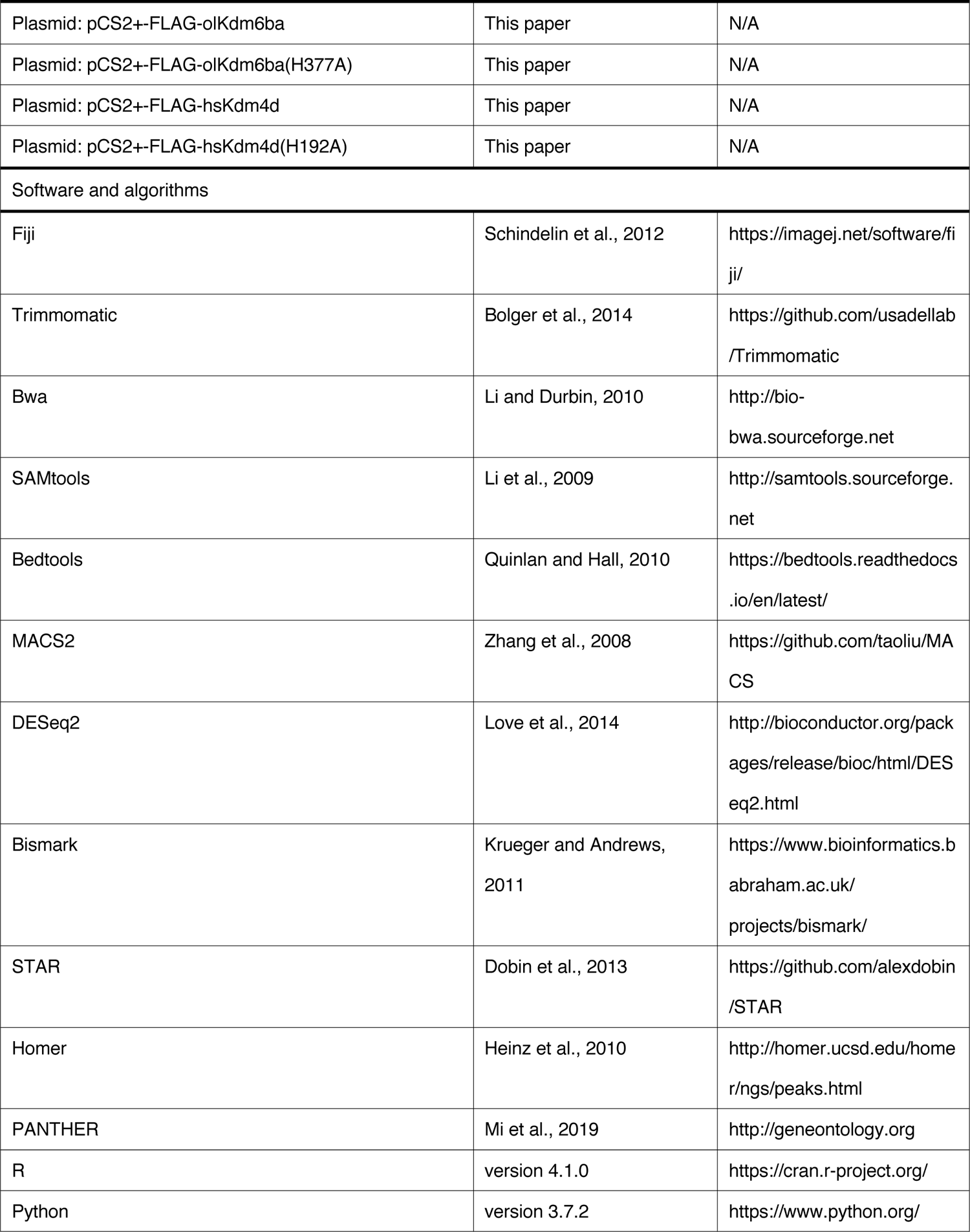

