## Supplementary figures for "Histone marks retained during epigenetic reprogramming and their roles essential for fish early development"


#### Authors/Affiliations

Hiroto S. Fukushima[1], Hiroyuki Takeda[1,*] and Ryohei Nakamura[1,2,*]

[1]Department of Biological Sciences, Graduate School of Science, The University of Tokyo, Tokyo, 113-0033, Japan

[2]Lead contact

### Supplementary Figures and Legends

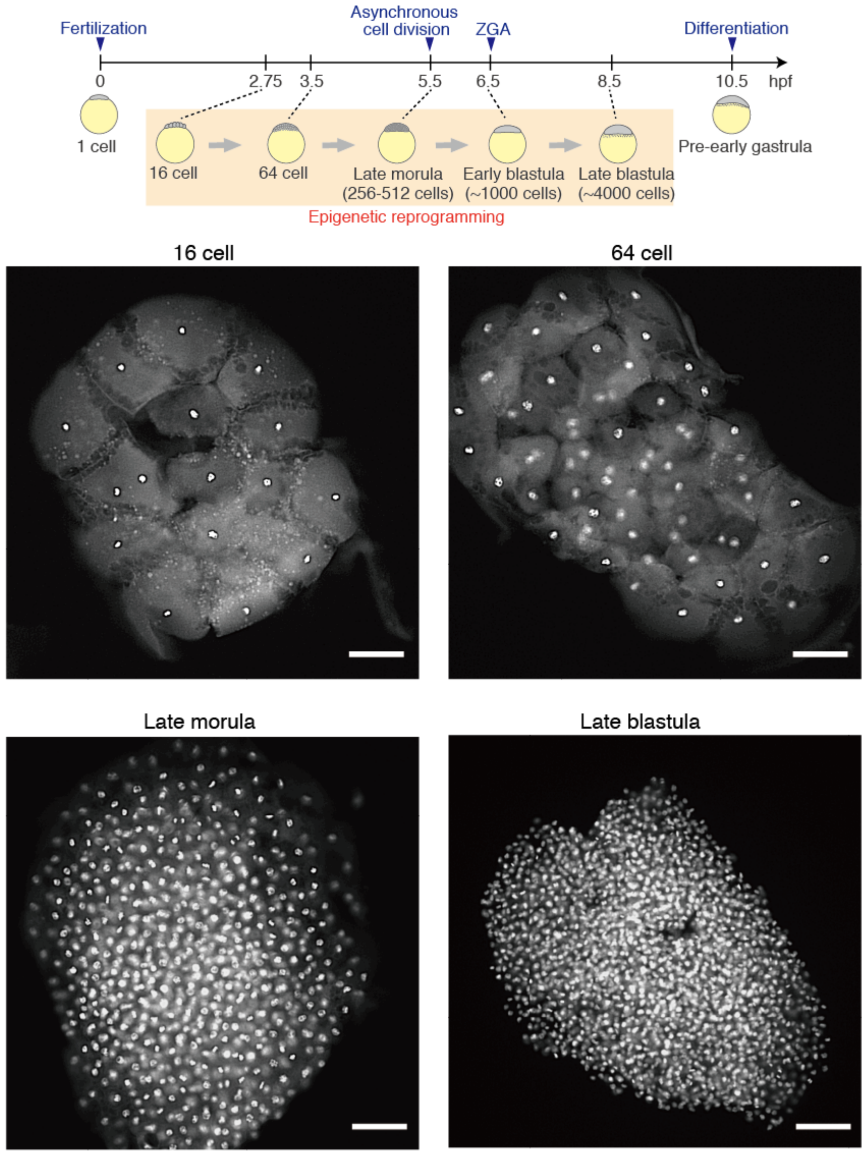


**Figure S1. Medaka development**
A schematic of medaka development and blastoderms after DAPI-staining at the four stages that we mainly focused on in this study. Scale bars indicate 100 $\mu$m.

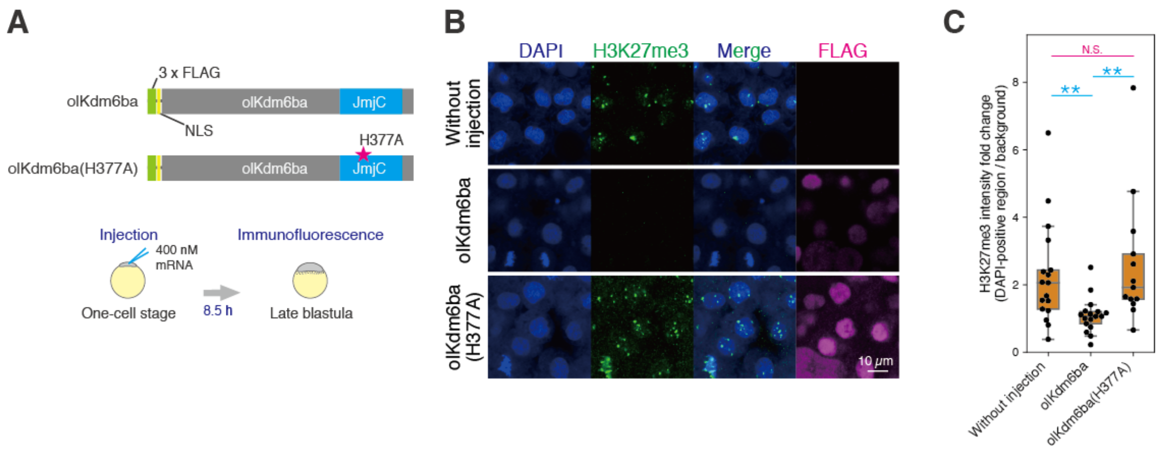


**Figure S2. Supporting data for Figure 1**

(A) Schematics of constructs (wildtype olKdm6ba and its catalytically inactive point mutant olKdm6ba(H377A)) and experimental design.

(B) Immunofluorescence staining at the late blastula stage showing depletion of H3K27me3 by olKdm6ba injection.

(C) Boxplots showing the signal intensities of each histone modification in DAPI-positive regions. Each dot indicates the average intensity in a single embryo. The intensity was normalized by background intensity. *** $p < 0.001$, ** $p < 0.01$, * $p < 0.5$, respectively.

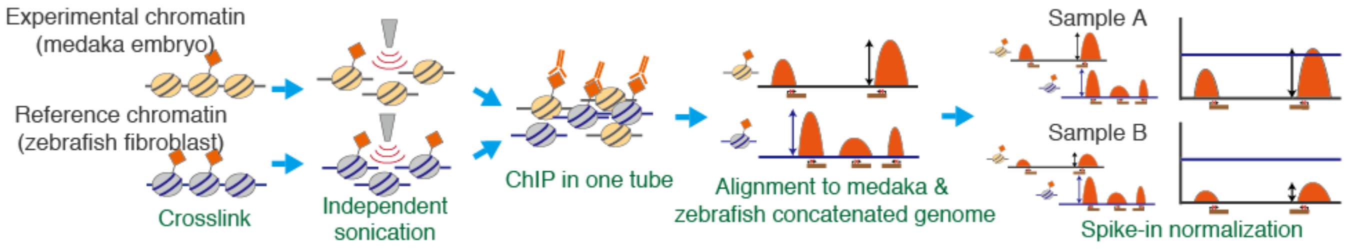


**Figure S3. A schematic illustration of spike-in ChIP-seq method used in this study**

We prepared reference chromatin from a zebrafish fibroblast cell line (BRF41) and mixed it with experimental chromatin (medaka embryo chromatin) in the same tube, which was then subjected to ChIP-seq. All reads were aligned to the medaka and zebrafish concatenated genome. Theoretically, the enrichment level of modifications in reference chromatin should be the same for all samples, so the level of histone modifications in experimental chromatin were normalized using the reference chromatin ChIP signal.

#### Figure S4

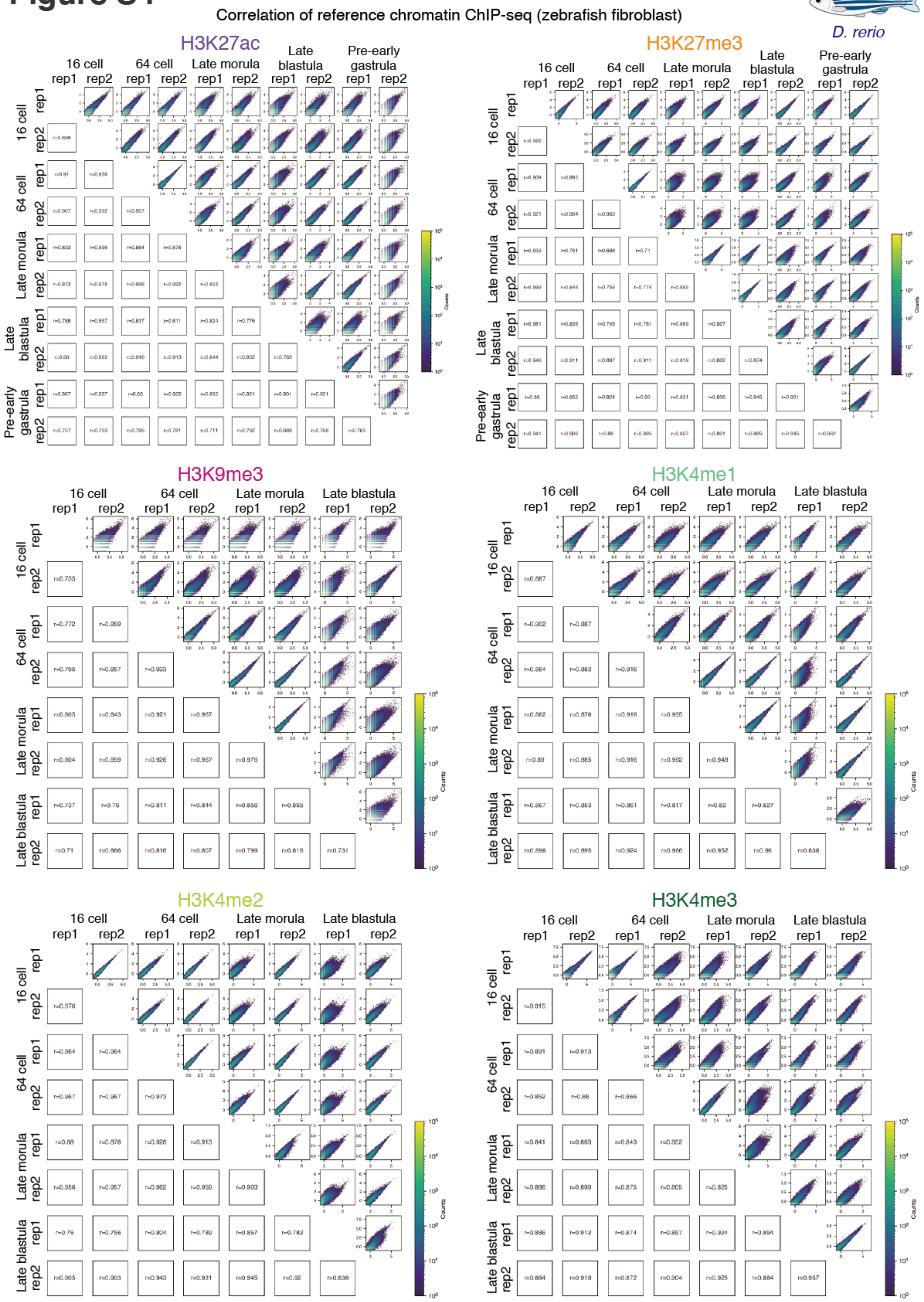


**Figure S4. Reproducibility of histone modification enrichment in reference chromatin (zebrafish fibroblast) revealed by spike-in ChIP-seq**

2D histograms showing correlation between each reference chromatin. Log2 (RPKMconv + 0.5) for each 5000 bp window-divided genomic interval and Pearson's correlation coefficients (r) are shown (RPKMconv: conventional RPKM).

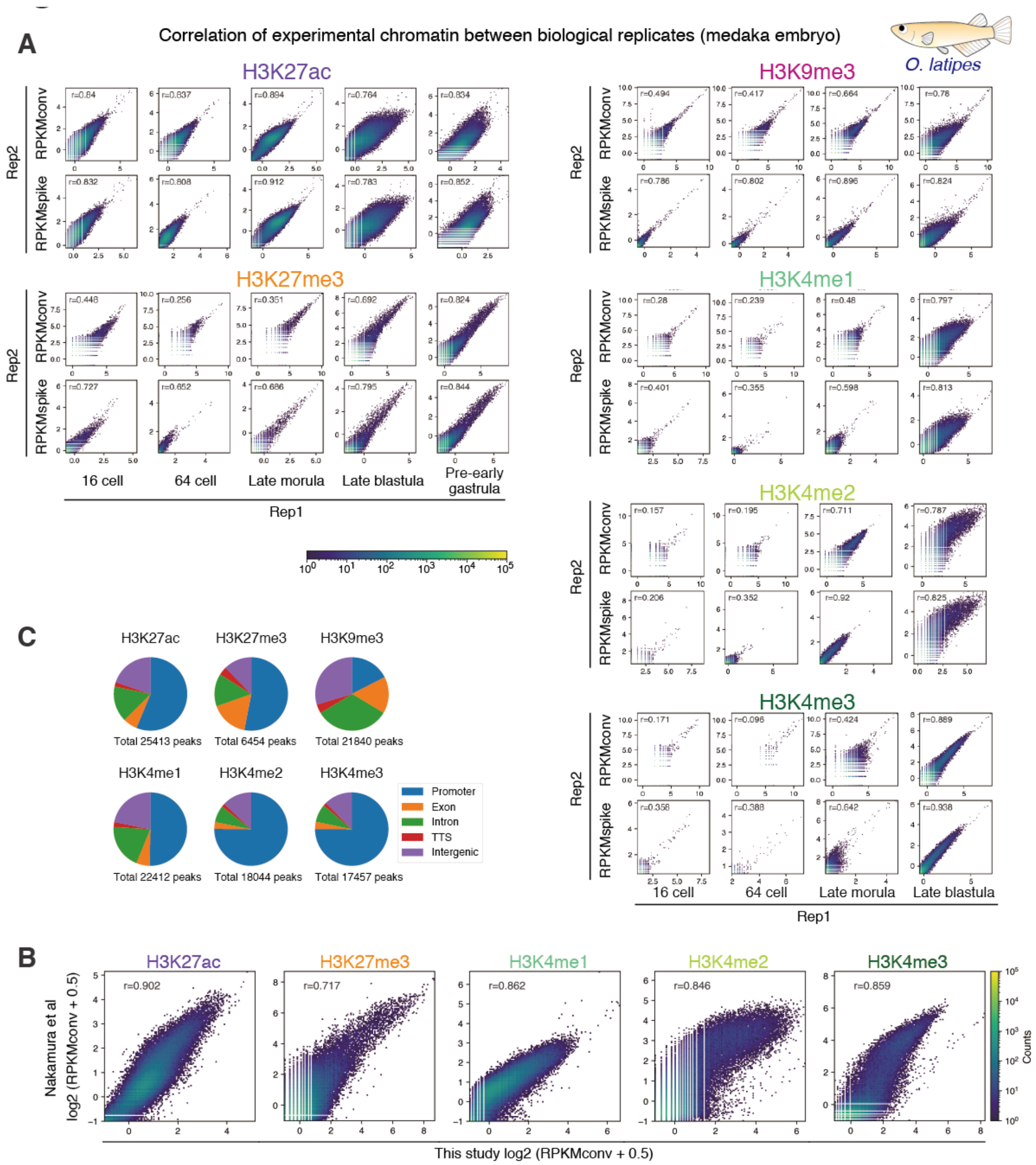


**Figure S5. Reproducibility of histone modification enrichment in experimental chromatin (medaka embryo) revealed by spike-in ChIP-seq**

(A) 2D histograms showing correlation between two biological replicates of experimental chromatin. Log2 (RPKMconv or RPKMspike + 0.5) for each 5000 bp window-divided genomic interval and Pearson's correlation coefficients (r) are shown (RPKMconv: conventional RPKM, RPKMspike: spike-in normalized RPKM).

(B) 2D histogram showing correlation between ChIP-seq data in this study and that in a previous study (Nakamura et al., 2014).

(C) Pie charts showing genomic distribution of spike-in ChIP-seq peaks for each modification.

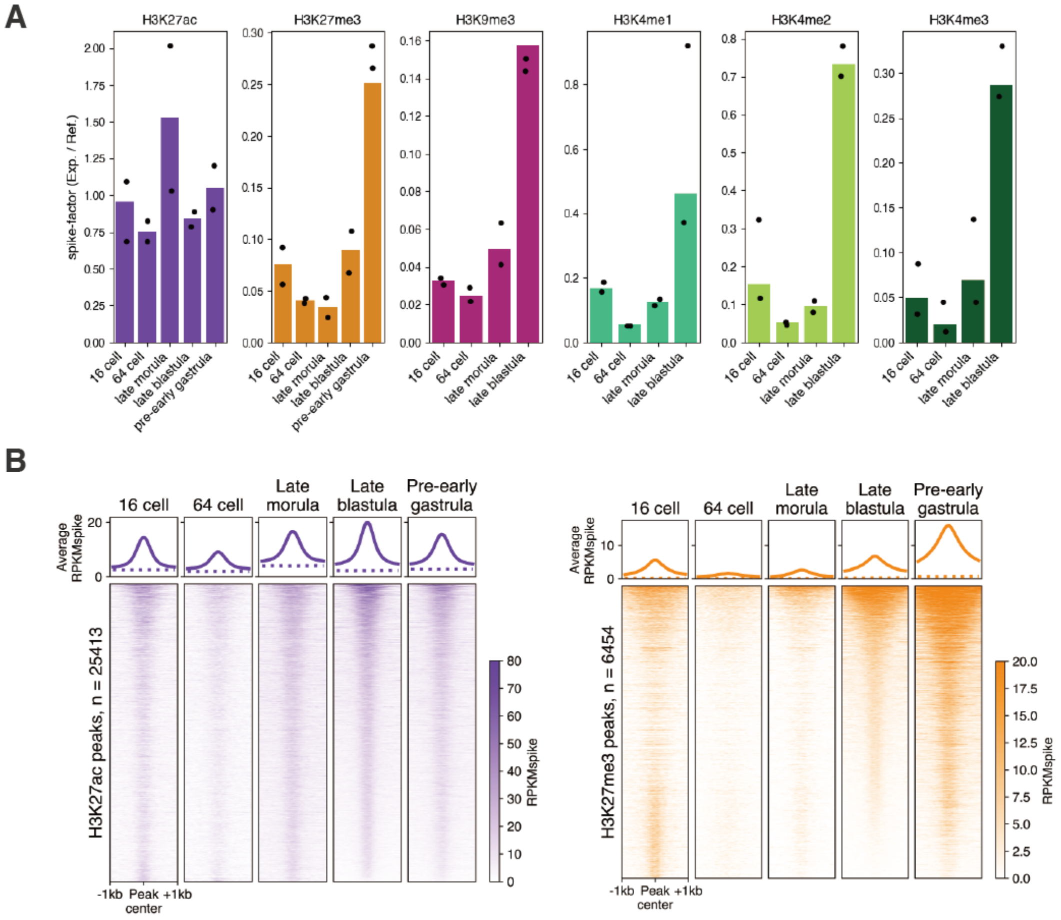


**Figure S6. Global level of each modification after spike-in normalization**
(A) Bar plots showing the global level of each modification at all stages calculated by spike-in ChIP-seq (see method for details). Here, bars and dots indicate values of pooled samples and individual replicates, respectively. We note that the values of pooled samples (bars) are not just the average of two replicates (dots) because deduplication affects the number of total reads.
(B) Genome-wide changes in enrichment of each modification including the pre-early gastrula stage data (also refer to Figure 2B). The average enrichment levels after spike-in normalization (RPKMspike) around all peaks and randomized peaks at each stage are shown as solid lines and dashed lines, respectively (top). Heatmaps showing enrichment levels (RPKMspike) around all peaks (± 1 kb from peak center).

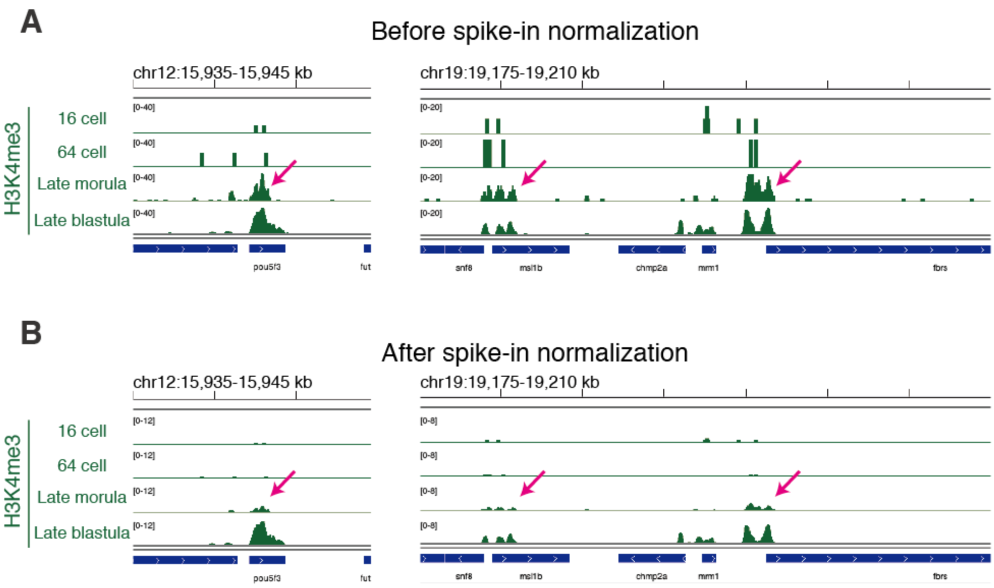


**Figure S7. Spike-in normalization revealed very limited accumulation of H3K4me3 at the late morula stage**
(A-B) Track views showing H3K4me3 enrichment before (A) and after spike-in normalization (B). Arrows indicate the genomic regions where H3K4me3 accumulation is observed before spike-in normalization.

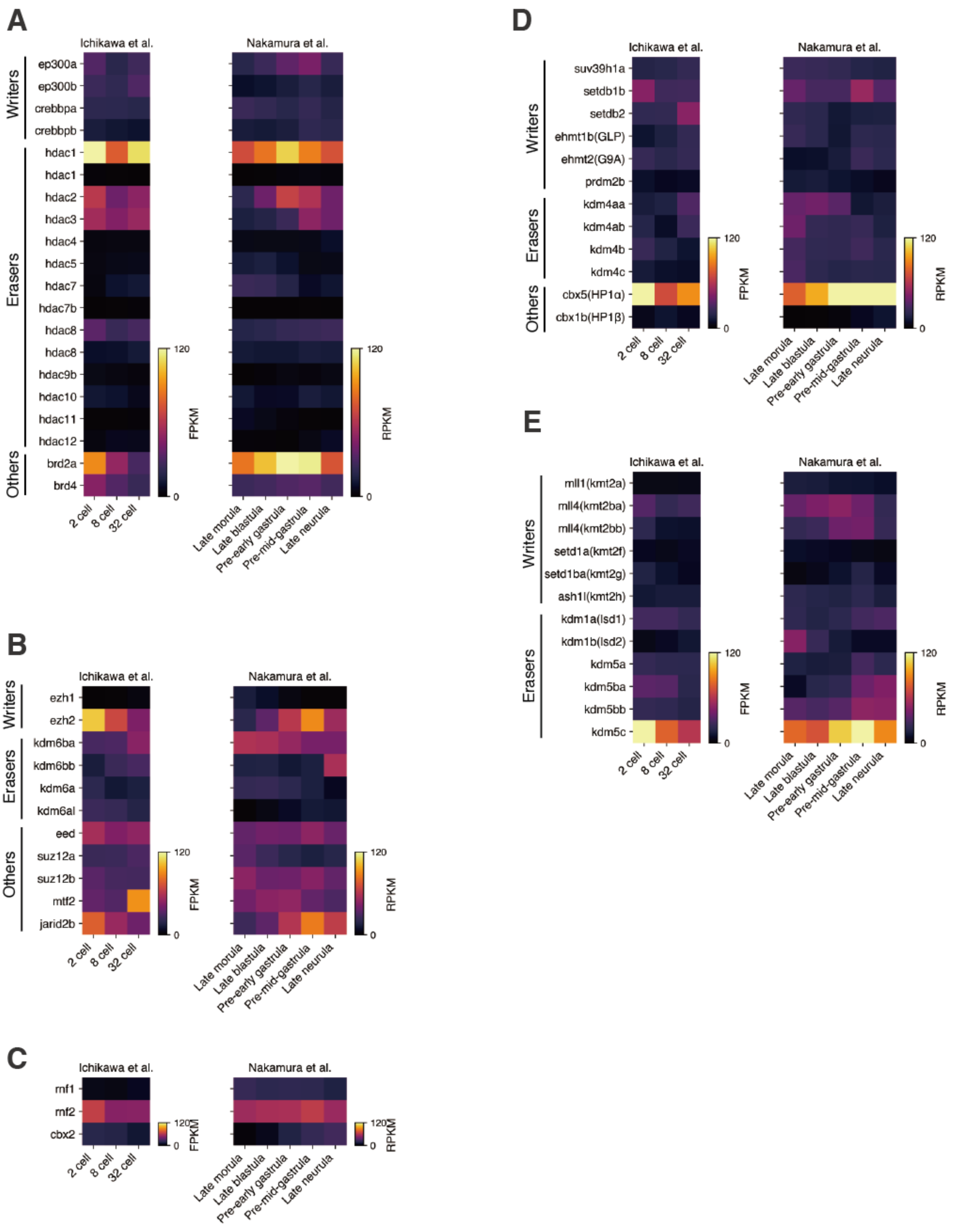


**Figure S8. Expression levels of writers, erasers and other related proteins of histone modifications**
(A) Expression levels of writers, erasers and readers of H3K27ac.
(B) Expression levels of writers, erasers and PRC2 components, related to H3K27me3.
(C) Expression level of PRC1 components, related to H3K27me3.
(D) Expression levels of writers, erasers and readers of H3K9me3.
(E) Expression levels of writers, erasers of H3K4 methylations.
RNA-seq data of the 2-cell, 8-cell and 32-cell stages were obtained from (Ichikawa et al., 2017) and the late morula, late blastula and pre-early gastrula stages were obtained from (Nakamura et al., 2021).

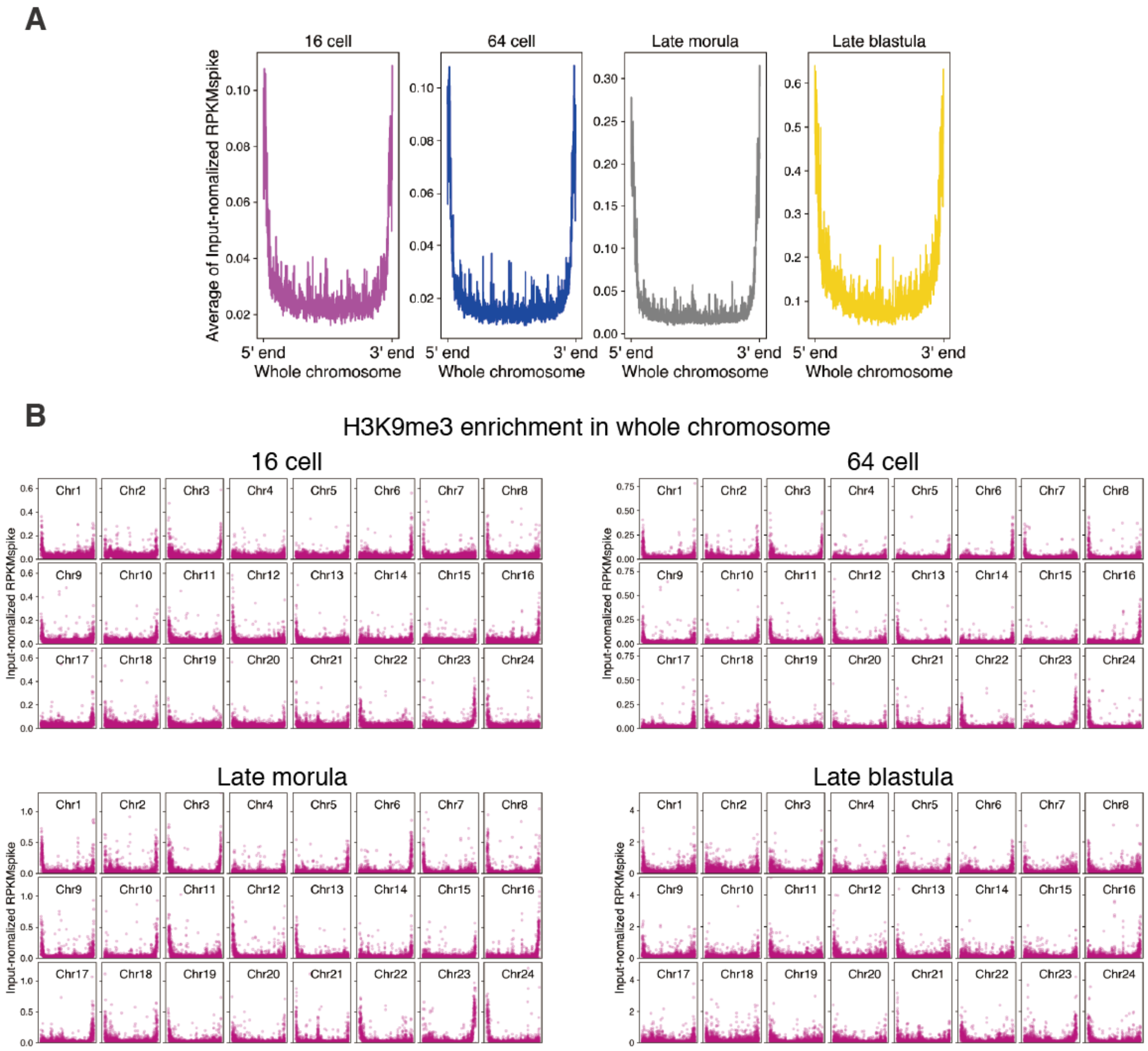


**Figure S9. Spike-in ChIP-seq showing retention of H3K9me3 localized at telomeric regions.**
(A) Average H3K9me3 signals of all chromosomes at four stages after spike-in normalization. The data in at each stage are shown separately here. To exclude repeat bias, the signal was further normalized by input signal (see Method).
(B) H3K9me3 enrichment along all individual chromosomes at four stages after spike-in normalization. Each dot represents input-normalized RPKMspike levels within 10 kb-divided genomic intervals.

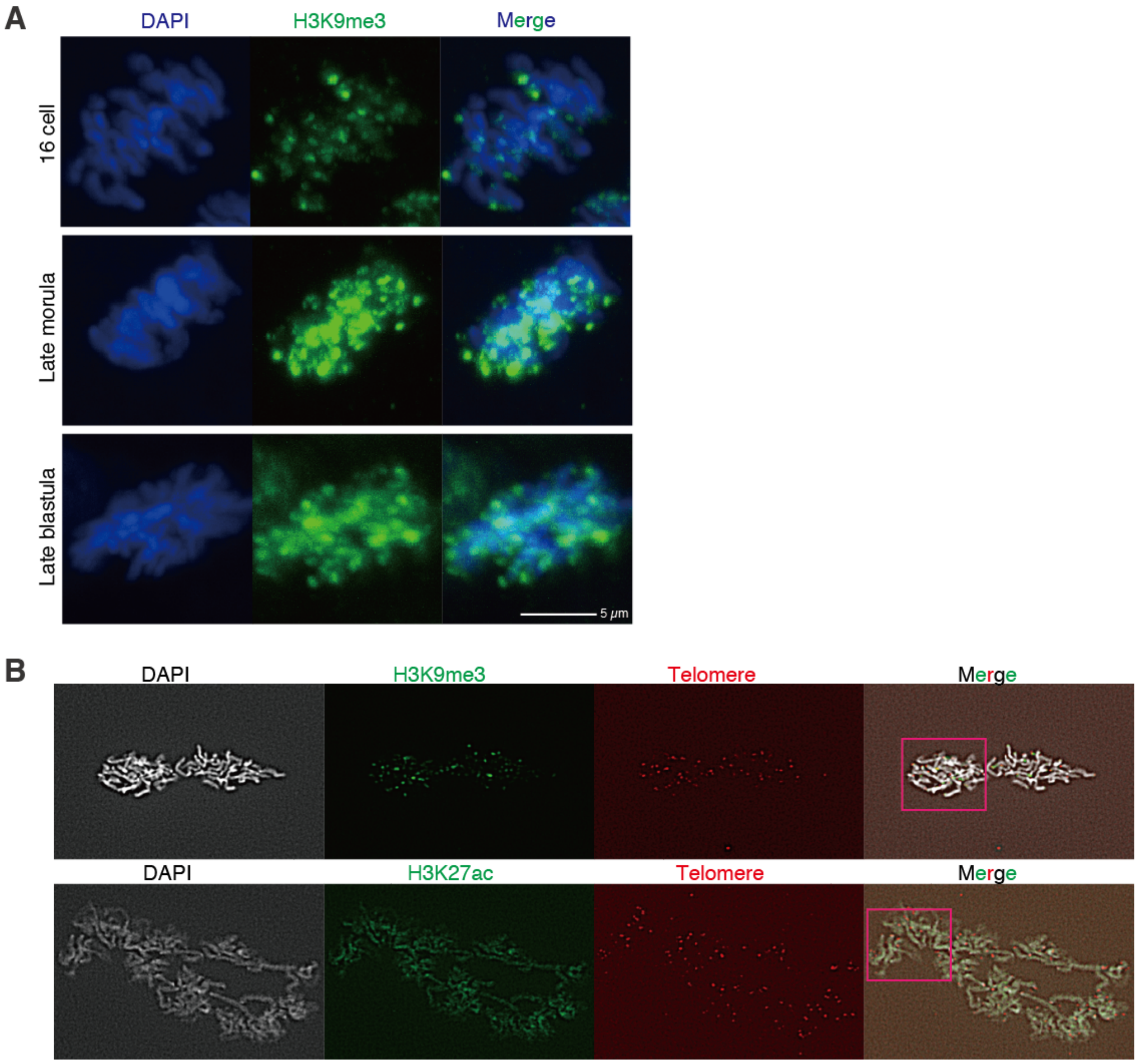


**Figure S10. Imaging data showing retention of H3K9me3 at telomeric regions.**
(A) Immunofluorescence staining of H3K9me3 in mitotic phase (metaphase or anaphase). See also Figure 3C (data at the 64-cell stage).
(B) Original data of Figure 3D. The areas marked in magenta are enlarged in Figure 3D.

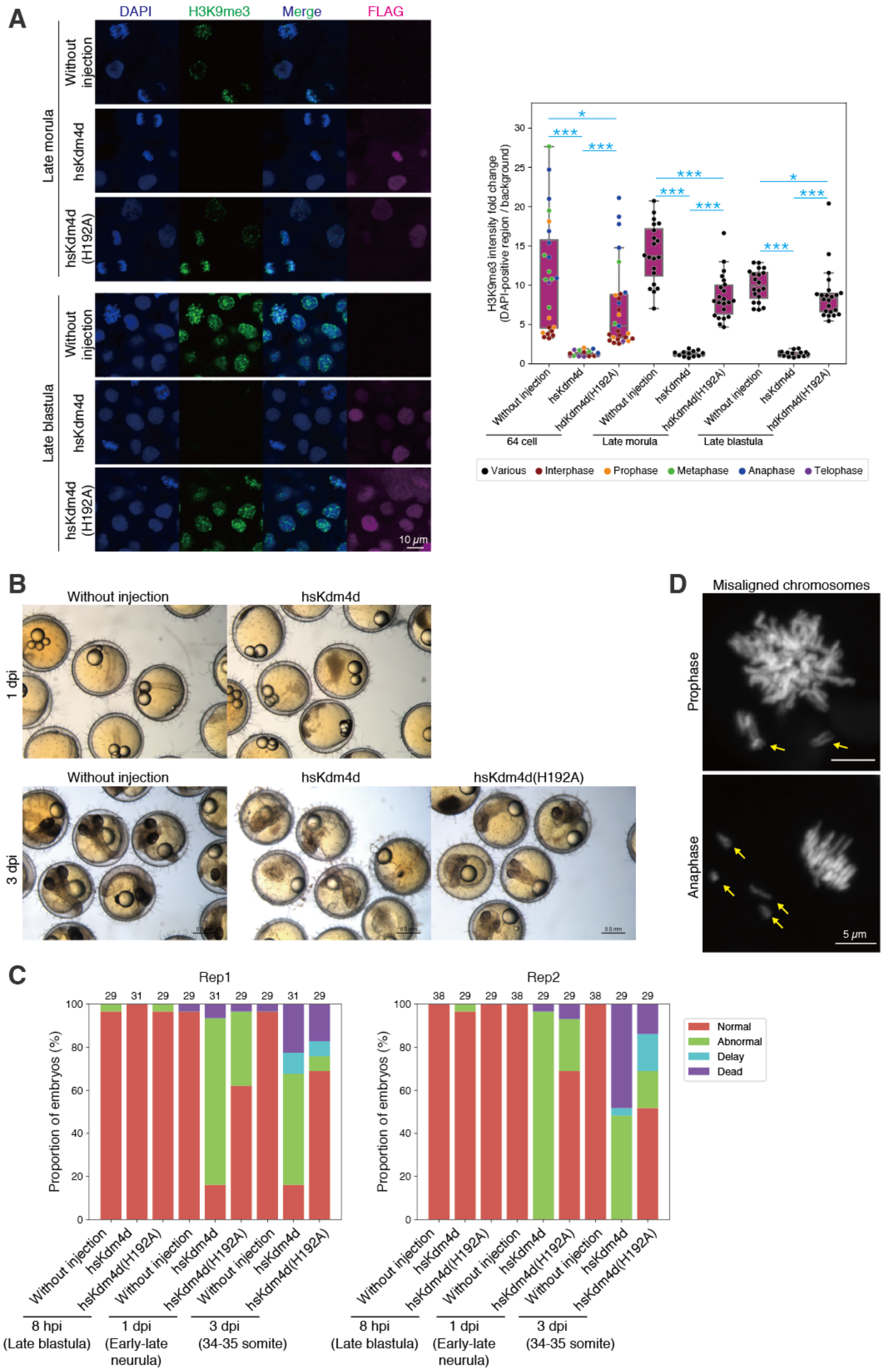


**Figure S11. Supporting data for hsKdm4d experiments**

(A) Immunofluorescence staining of H3K9me3 and FLAG (left), and boxplots showing the signal intensities of each histone modification in DAPI-positive regions (right). Each dot indicates the average intensity in a single embryo. FLAG signal indicates the enrichment level of translated hsKdm4d proteins. The intensity was normalized by background intensity. Cell cycle phases at the 64-cell stage embryos are shown as dots with different colors. Phases of the cell cycle after the late morula stage are not indicated because cells divide asynchronously from the late morula stage. See also Figure 4A (schematics of experiments) and 4B (data at the 64-cell stage). *** $p < 0.001$, ** $p < 0.01$, * $p < 0.5$, respectively.

(B) Phenotypes of embryos injected with hsKdm4d.

(C) Percentages of embryos showing each phenotype. Two biological replicates are shown separately. The number above each bar indicates the total number of embryos in each condition. *** $p < 0.001$, ** $p < 0.01$, * $p < 0.5$, respectively. hpi/dpi = hours/days post injection.

(D) Representative phenotypes of chromosome segregation errors (misaligned chromosomes). Arrows indicate the errors. See also Figure 4C and 4D.

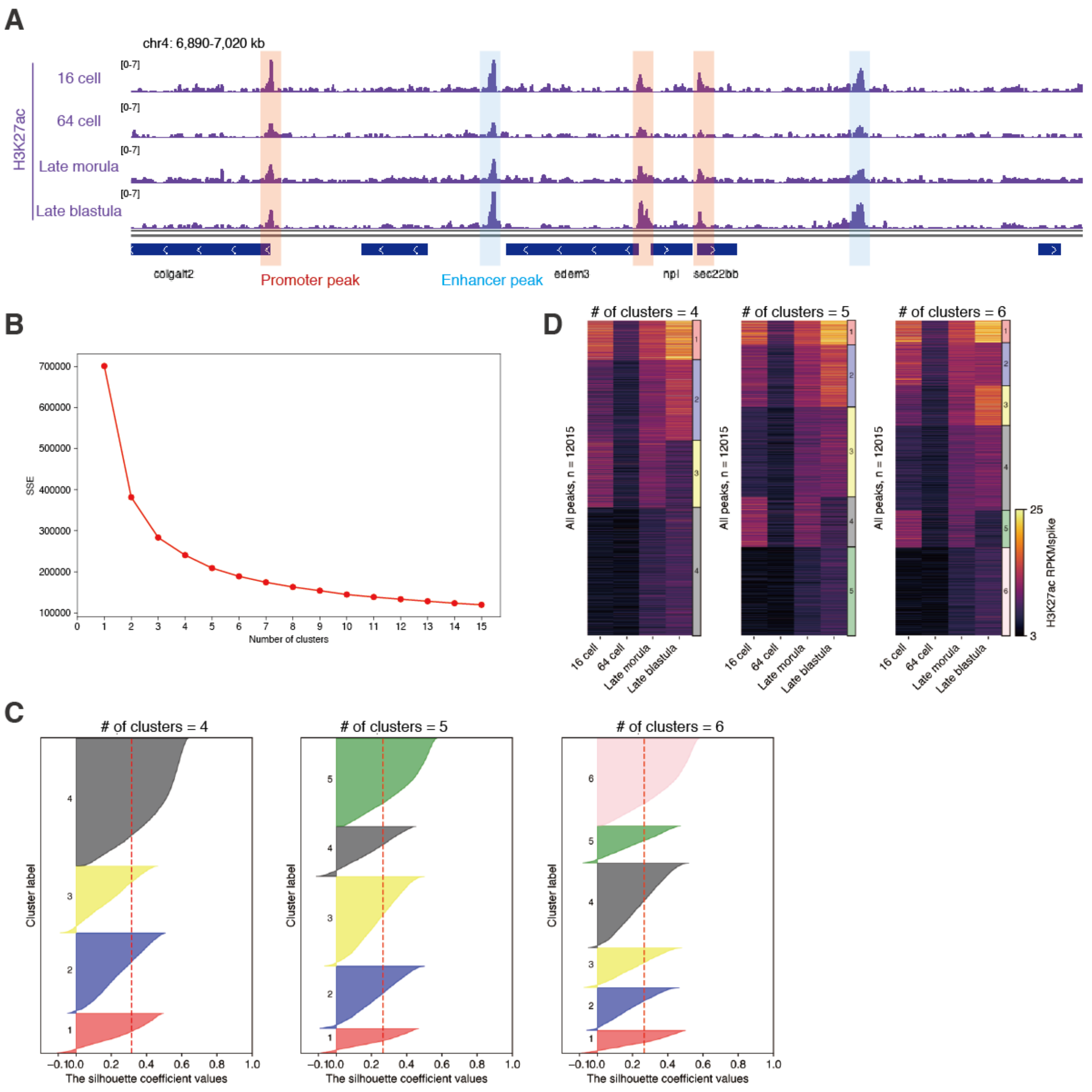


**Figure S12. Decision of number of clusters for k-means clustering of H3K27ac-marked promoters.**
(A) Track view showing promoter peaks (peaks within -2.5 kb ≤ TSS ≤ 2.5 kb, red) and enhancer peaks (peaks at least 2 kb away from promoter peaks, blue) of H3K27ac.
(B) Elbow method for the decision of optimal cluster number for k-means. Line plot showing sum of squared errors (SSE) after k-means clustering titrating the number of clusters. This analysis showed that the SSE starts decreasing in an almost linear fashion from cluster number = 5, suggesting that the optimal number of clusters is around 5.
(C) Silhouette analysis for the deciding of optimal cluster number for k-means. Horizontal bar plots showing the silhouette coefficient value in each element, and dashed line indicates the average silhouette coefficient. Ideally, each cluster should show as many as possible elements with a higher silhouette coefficient. From this perspective, we concluded that cluster number 5 or 6 is better than 4.
(D) A heatmap showing H3K27ac enrichment in H3K27ac-marked promoters after k-means clustering using different cluster numbers. Based on the results of Figure S12A, the heatmaps using k = 4, 5 or 6 were compared. However, the cluster showed similar characters between those heatmaps. Therefore, we concluded that the best number of clusters for k-means was 5.

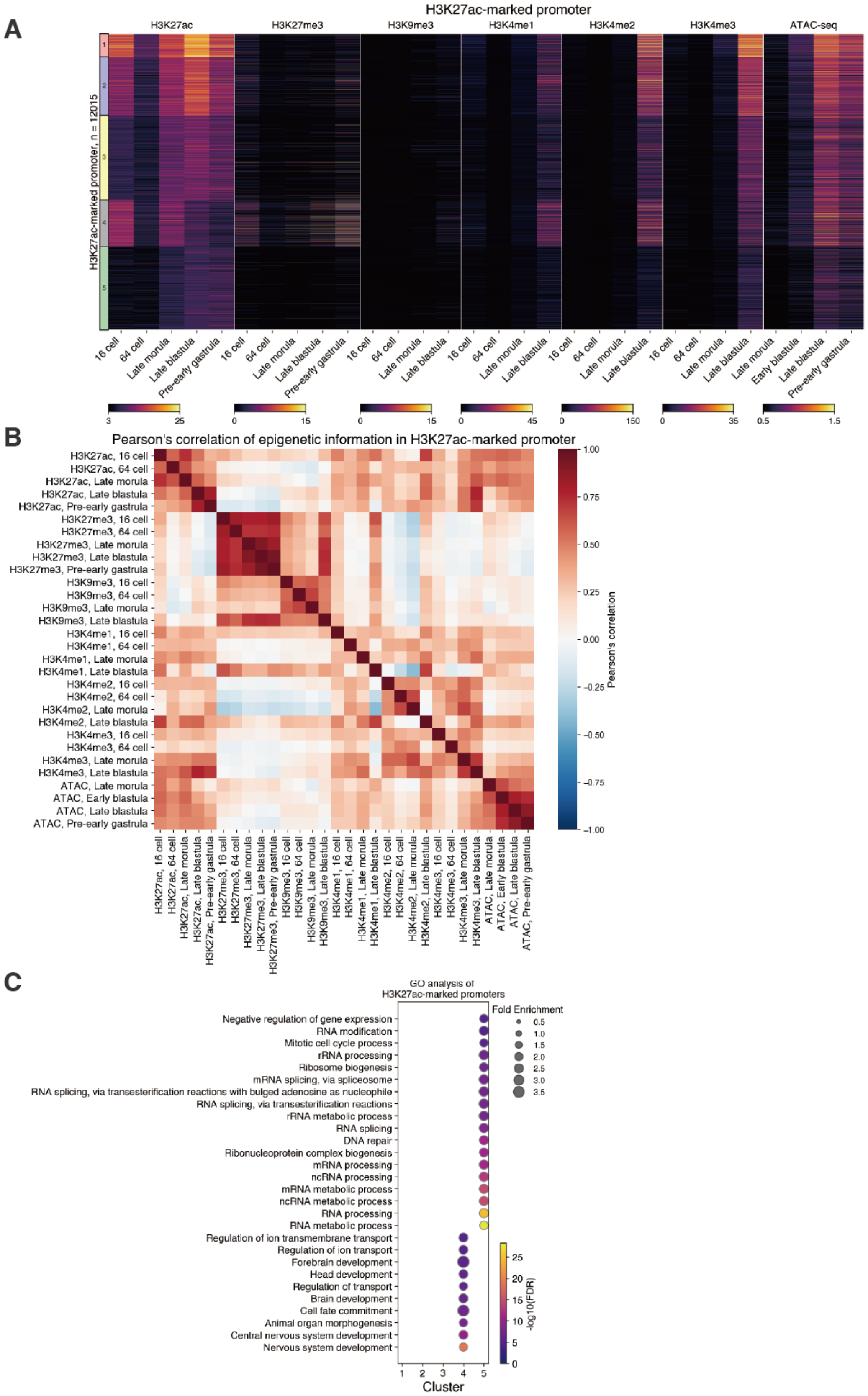


**Figure S13. Characters of H3K27ac-marked promoters.**

(A) Heatmaps showing epigenetic modification levels at H3K27ac-marked promoters.

(B) A heatmap showing Pearson's correlation of epigenetic information in H3K27ac-marked promoters.

(C) GO terms (biological process) enriched in the genes associated with each H3K27ac-marked promoter cluster. Only those GO terms (biological process) which have an FDR less than $10^{-5}$ and whose enrichment more than 2.1 fold were listed. Color indicates FDR, and circle size is the fold enrichment of each GO term.

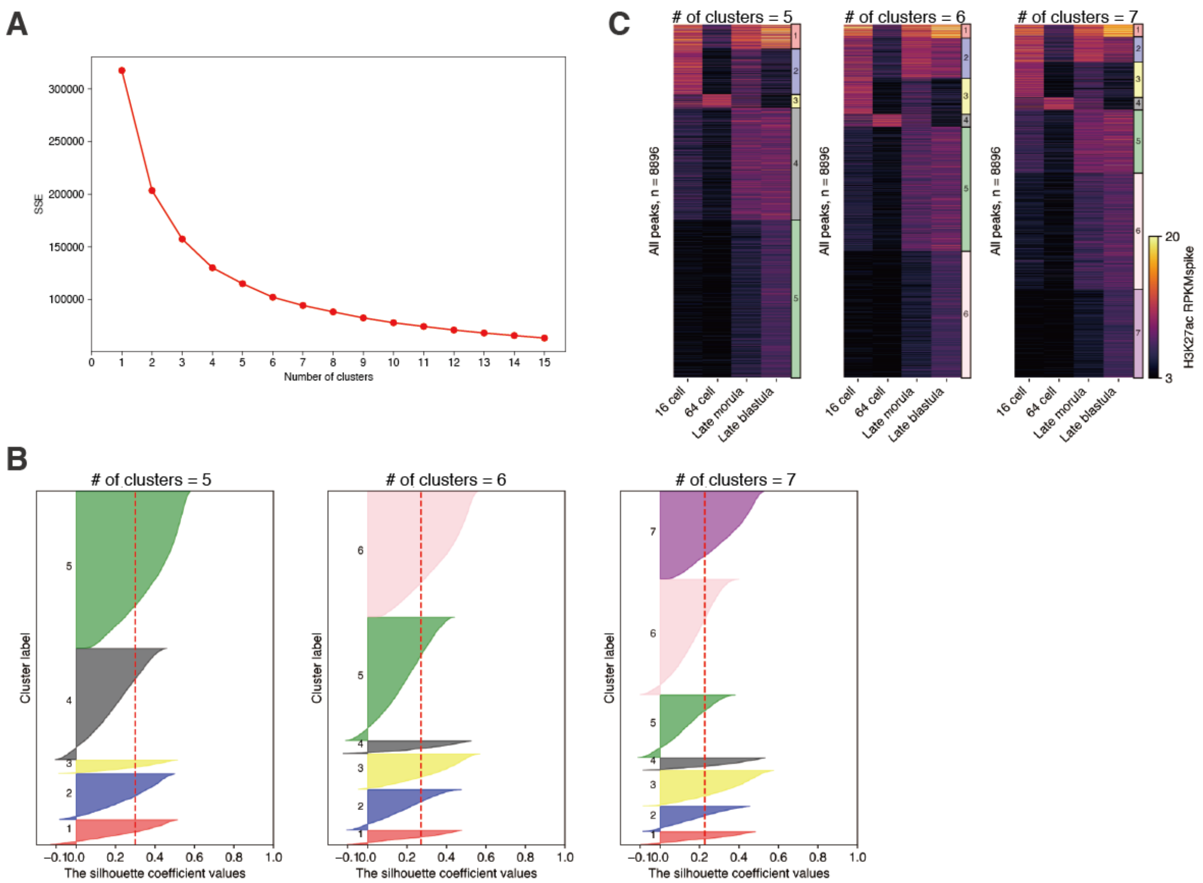


**Figure S14. Decision of number of clusters for k-means clustering of H3K27ac-marked enhancers.** (A) Elbow method for the decision of optimal cluster number for k-means. Line plot showing sum of squared errors (SSE) after k-means clustering titrating the number of clusters. From this, the SSE starts decreasing in an almost linear fashion from cluster number = 6, suggesting that the optimal number of clusters is around 6.
(B) Silhouette analysis for deciding of optimal cluster number for k-means. Horizontal bar plots showing silhouette coefficient value in each element, and dashed line indicating the average silhouette coefficient. Ideally, each cluster should show as many as possible elements with higher silhouette coefficient. From this perspective, we concluded that cluster number 6 is optimal.
(C) Heatmaps showing H3K27ac enrichment in H3K27ac-marked promoters after k-means clustering using different cluster numbers. Based on the results of Figure S14A, the heatmaps using several numbers of clusters = 5, 6 or 7 were compared. Given that the cluster having very different character was not found among those heatmaps, we concluded that the best number of clusters for k-means was 6.

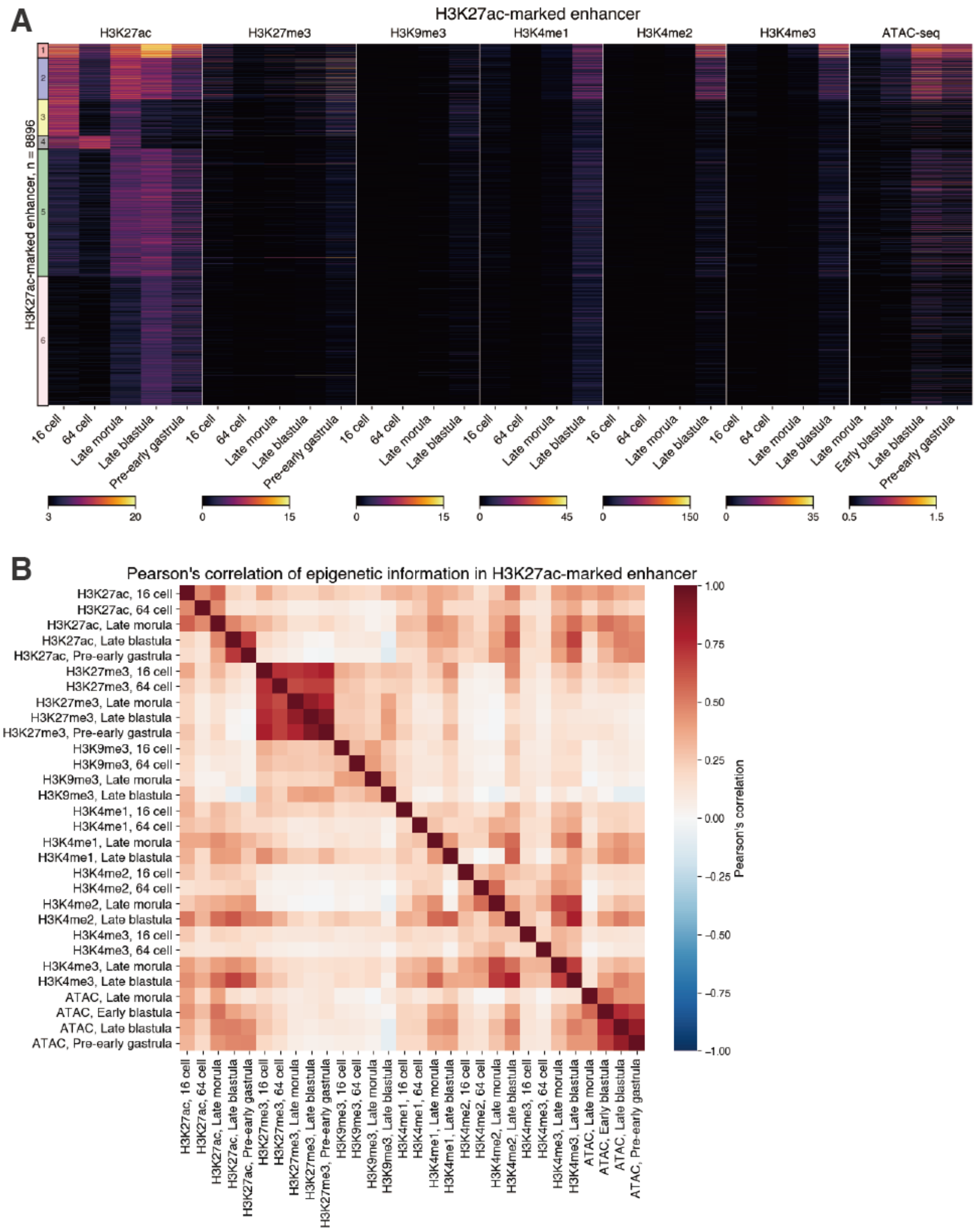


**Figure S15. Characters of H3K27ac-marked enhancers**

(A) A heatmap showing epigenetic modifications of H3K27ac-marked enhancers.

(B) A heatmap showing Pearson's correlation of epigenetic information in H3K27ac-marked enhancers.

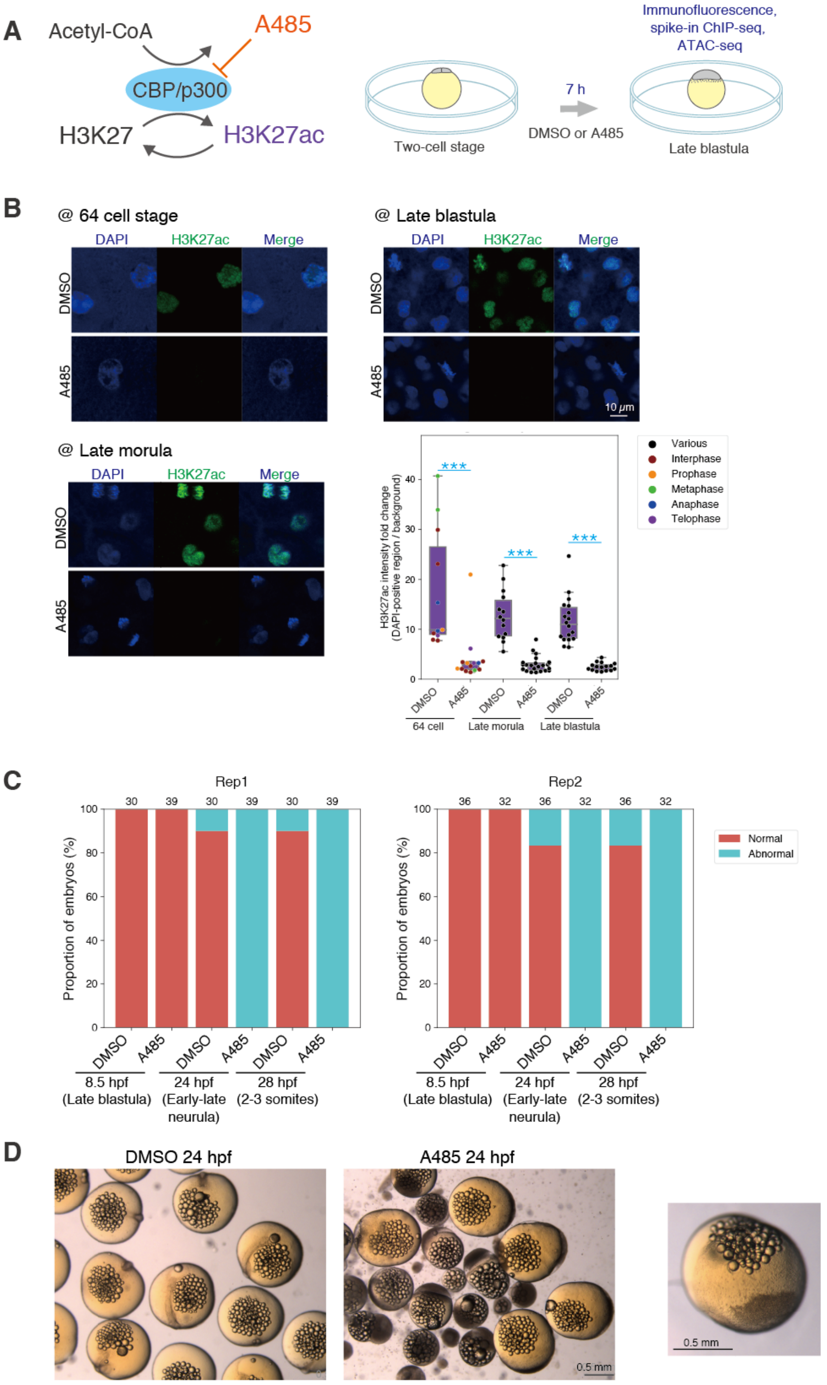


**Figure S16. Phenotype and global change in H3K27ac accumulation after A485 treatment.**
(A) Schematics of A485 experiments. CBP/p300 induces H3K27ac, and acetyl-CoA is its substrate. A485 competitively inhibits CBP/p300 catalytic activity (Lasko et al., 2017). Dechorionated embryos were incubated with DMSO or 20 $\mu$M A485 from the two-cell stage. After 7 hours of incubation, the embryos were used for immunofluorescence staining, ChIP-seq and ATAC-seq.
(B) Immunofluorescence staining of H3K27ac (left) and boxplots showing the signal intensities of each histone modification in DAPI-positive regions (right). Each dot indicates the average intensity in a single embryo. The intensity was normalized by background intensity. Cell cycle phases at the 64-cell stage embryos are shown as dots with different colors. Phases of cell cycle after the late morula stage are not indicated because cells divide asynchronously from the late morula stage. *** $p < 0.001$, ** $p < 0.01$, * $p < 0.5$, respectively.
(C) Percentages of normal and abnormal embryos. Two biological replicates are shown separately. The number above each bar indicates the number of total embryos in each condition.
(D) Phenotypes observed at 24 hpf. Embryonic body was normally formed in DMSO-treated embryos while gastrulation arrest was observed in A485-treated embryos. The representative image of A485-treated embryos was enlarged in the panel on the right. hpf = hours post fertilization.

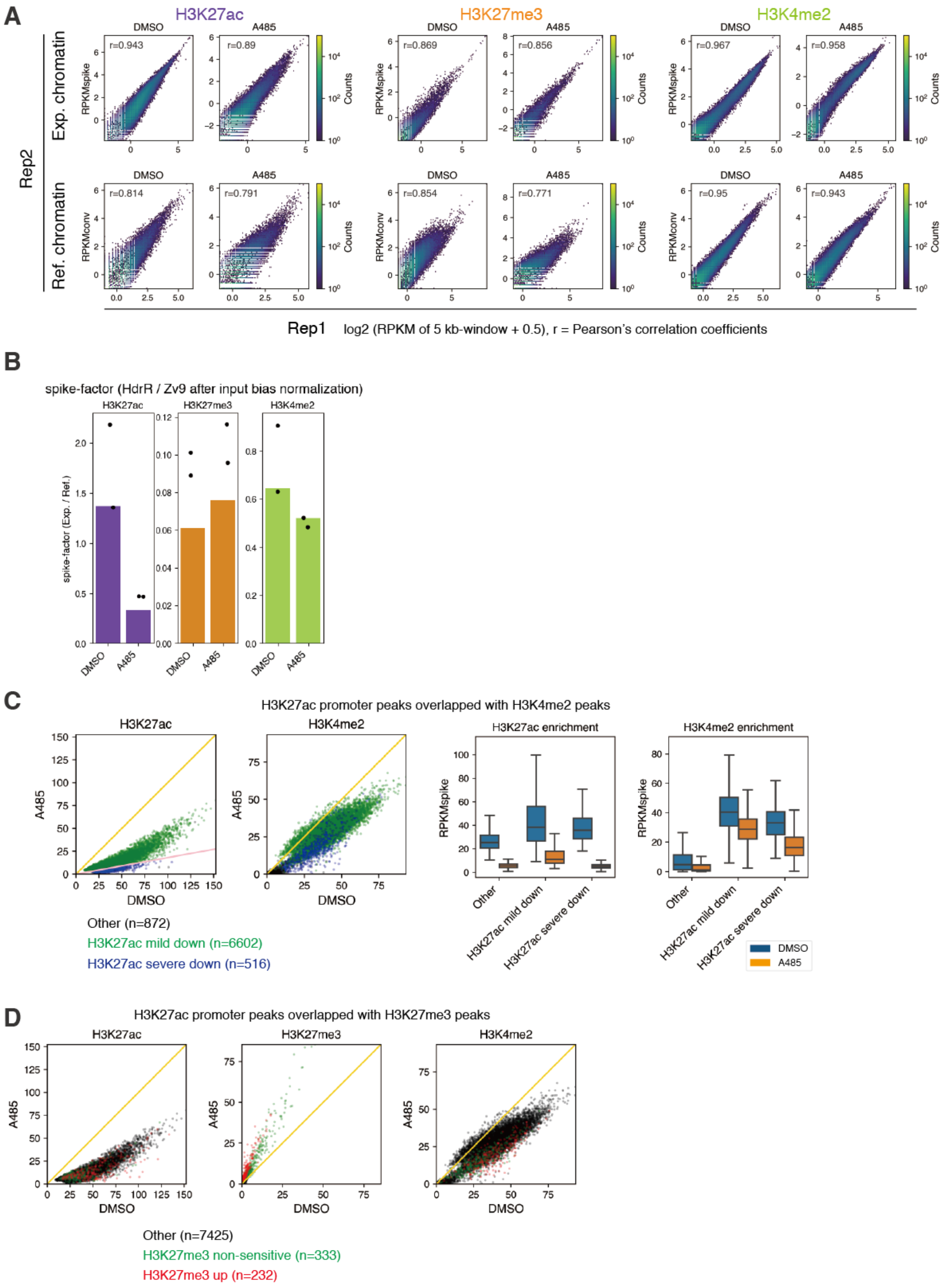


**Figure S17. Analysis of spike-in ChIP-seq data of A485-treated embryos.**
(A) 2D histograms showing correlation between two biological replicates of experimental and reference chromatin. Log2 (RKPMspike or RPKMconv + 0.5) for each 5000 bp window-divided genomic interval and Pearson's correlation coefficients (r) are shown (RPKMconv: conventional RPKM, RPKMspike: spike-in normalized RPKM).
(B) Bar plots showing the global level of each modification at all stages calculated by spike-in ChIP-seq (see method in detail). Here, bars and dots indicate values of pooled samples and individual replicates, respectively. We note that the values of pooled samples (bars) are not just the average of two replicates (dots) because deduplication after pooling of two replicates affects the number of reads (see method).
(C) Comparison of H3K27ac and H3K4me2 enrichment in H3K27ac peaks between DMSO and A485 treatment by scatter plots (left) and boxplots (right). "Other" indicates the H3K27ac peaks without H3K4me2 enrichment. The yellow line and pink line in the scatter plot indicate y=x and y=0.18*x (threshold for "severe down"), respectively.
(D) Scatter plot showing H3K27ac and H3K27me3 enrichment in H3K27ac peaks in DMSO and A485 treatment. "Other" indicates the H3K27ac peaks without H3K27me3 enrichment. Yellow line indicates y=x.

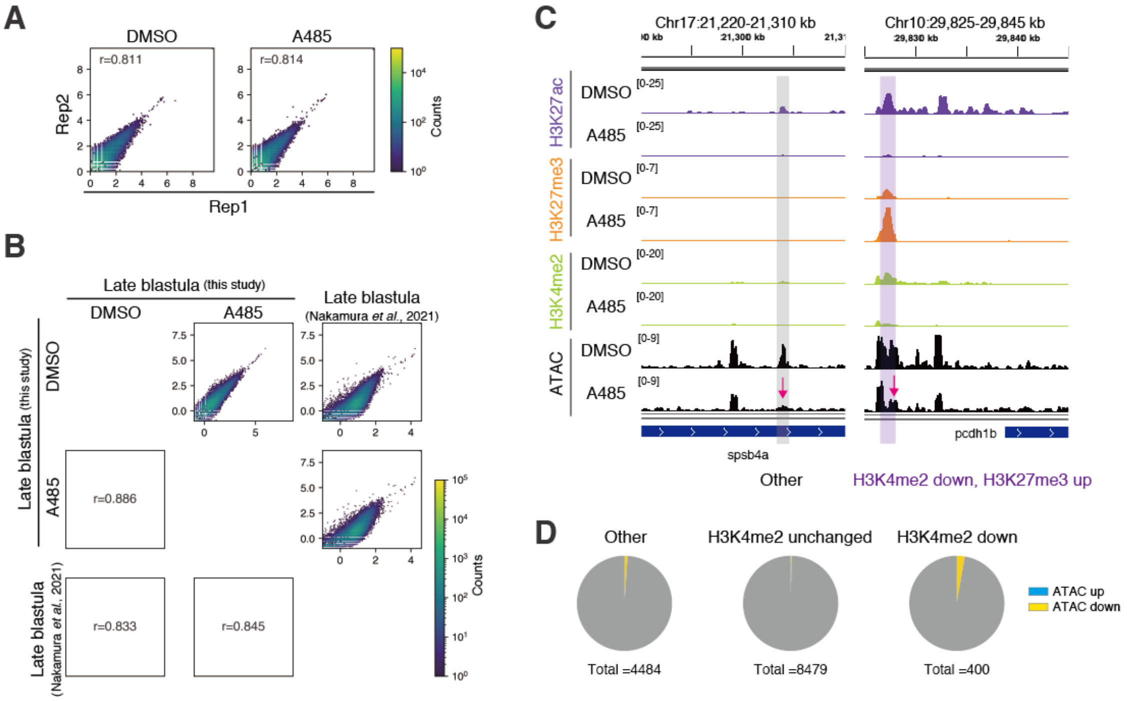


**Figure S18. Analysis of ATAC-seq data of A485-treated embryos.**
(A) 2D histograms showing correlation between two biological replicates of experimental and reference chromatin. Log2 (RPKMconv + 0.5) for each 5000 bp window-divided genomic interval and Pearson's correlation coefficients (r) are shown (RPKMconv: conventional RPKM).
(B) 2D histograms showing correlation between experiment in this study and data from a previous study (Nakamura et al., 2021). RPKMconv of pooled samples are used here. Log2 (RPKMconv + 0.5) for each 5000 bp window-divided genomic interval and Pearson's correlation coefficients (r) are shown (RPKMconv: conventional RPKM).
(C) Track views showing the histone modification enrichments after spike-in normalization and chromatin accessibility in DMSO or A485-treated embryos. The magenta arrow indicates where chromatin accessibility was altered after A485 treatment. See also in Figure 6A.
(D) Pie charts showing the proportion of H3K27ac peaks whose chromatin accessibility was reproducibly affected by A485 treatment.

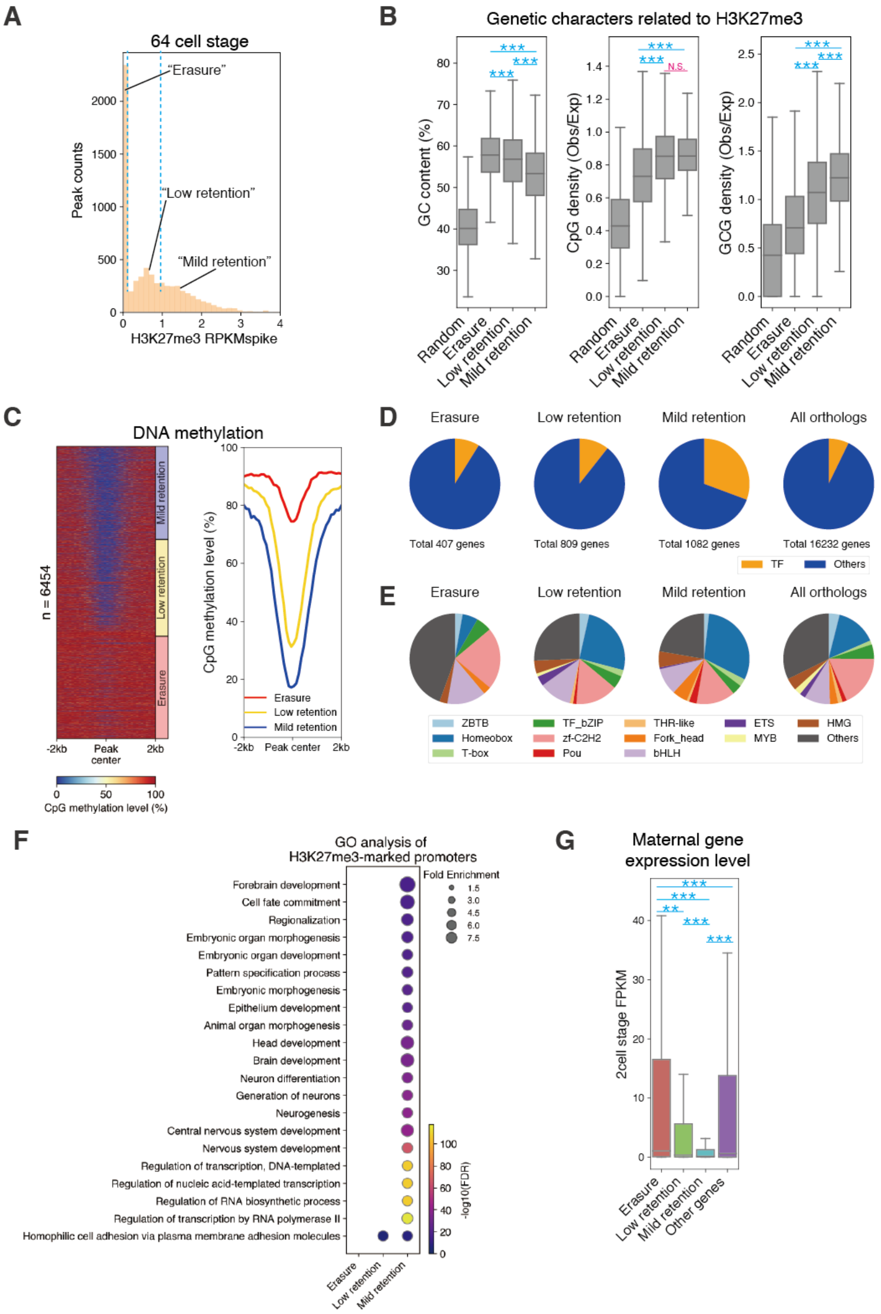


**Figure S19. Genomic regions silenced in oocytes and having high-affinity for Polycomb residually retain H3K27me3.**

(A) A histogram of H3K27me3 levels after spike-in normalization (RPKMspike) in H3K27me3 merged peaks at 64 cell stage. As indicated with dashed lines, peaks were divided into three groups based on H3K27me3 RPKMspike value at the 64-cell stage (“Erasure”: RPKMspike = 0, “Low retention”: 0 < RPKMspike ≤ 1, “Mild retention”: 1 < RPKMspike).
(B) Boxplots showing enrichment of genetic characters in each group. *** p < 0.001, ** p < 0.01, * p < 0.5, respectively.
(C) A heatmap showing DNA methylation levels around all H3K27me3 peaks (± 2 kb from peak center, window = 100 bp) (left) and line plot showing their averages (right). The order was sorted by RPKMspike of H3K27me3 at the 64-cell stage in heatmap. DNA methylation data was obtained from (Qu et al., 2012).
(D-E) Pie charts indicating the distribution of transcription factors (TFs) in each group (D) and pie charts showing the percentage of transcription factor family in each H3K27me3 peak group (E). Only those H3K27me3 peaks which are located at promoters and whose downstream gene have a human ortholog were included in this analysis. If several peaks are included in same promoter, the representative peaks which have highest RPKMspike value at the 64-cell stage were used. Human orthologs and human transcription factor data were obtained from Ensembl and AnimalTFDB 3.0 (Hu et al., 2019), respectively.
(F) GO terms (biological process) enriched in genes associated with H3K27me3-marked promoters. Only those GO terms with FDR less than $10^{-5}$ and with an enrichment of more than 3.3 fold are listed. Color indicates FDR, and circle size correlates to the fold enrichment of each GO term.
(G) Boxplots showing gene expression levels at the two-cell stage (Ichikawa et al., 2017) of genes in each group.

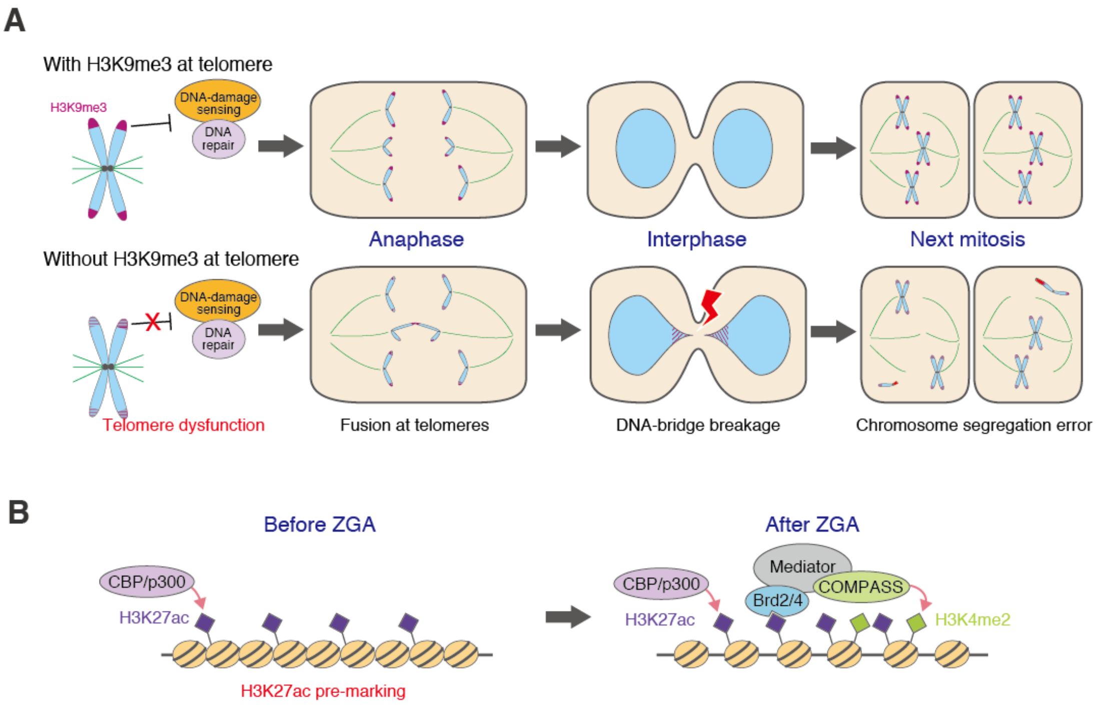


**Figure S20. Models.**
(A) Schematics of residual retention of H3K9me3 at telomeric regions at early developmental stages. During early development, H3K9me3 accumulation is mainly limited to telomeric regions. Without H3K9me3 at telomeric regions, telomere dysfunction is induced and results in fusion of two chromosomes and subsequent DNA-bridge breakage, suggesting that H3K9me3 prevents telomeres from recruiting DNA-damage sensing and/or DNA repair machinery.
(B) Schematics of H3K27ac pre-marking. CBP/p300 induces H3K27ac continuously after fertilization. After ZGA, H3K27ac reader BRD2/4 or recruitment of cofactors such as mediator and COMPASS may induce H3K4me2. These genomic regions become open after ZGA, but H3K27ac is not required for this gain of chromatin accessibility.
